# Lyl-1 regulates primitive macrophages and microglia development

**DOI:** 10.1101/2020.09.28.316570

**Authors:** Shoutang Wang, Deshan Ren, Anna-Lila Kaushik, Gabriel Matherat, Yann Lécluse, Dominik Filipp, William Vainchenker, Hana Raslova, Isabelle Plo, Isabelle Godin

**Author notes:** ***Corresponding author:*** Isabelle Godin, INSERM U1287; Institut Gustave Roussy-PR1; 114, rue Edouard Vaillant; 94805 VILLEJUIF Cedex; France, Email address. Department of Pathology and Immunology, Washington University School of Medicine, St. Louis, MO, 63110, USA. Medical school of Nanjing university, Model Animal Research Center, Nanjing University, Nanjing 210093, China (Deshan Ren). Plasseraud IP, 33064 Bordeaux, France. Agence Nationale pour la Recherche, Paris, France (Gabriel Matherat). Equal contribution (Shoutang Wang, Deshan Ren).

## Abstract

During ontogeny, macrophages (MΦ) populations emerge in the Yolk Sac (YS) via two distinct progenitor waves, prior to hematopoietic stem cell development. MΦ-progenitors from the primitive/”early EMP” and transient-definitive/”late EMP” waves both contribute to various resident-MΦ populations in the developing embryonic organs. Identifying factors that modulates early stages of MΦ-progenitor development may lead to a better understanding of defective function of specific resident-MΦ subsets.

Here we show that primitive macrophage (MΦ^Prim^) progenitors in the YS express Lyl-1, a bHLH transcription factor related to SCL/Tal-1. Transcriptomic analysis of YS MΦ-progenitors indicated that MΦ^Prim^ progenitors present at embryonic day (E) 9 are clearly distinct from those present at later stages. Disruption of Lyl-1 basic helix-loop-helix domain led initially to an early increased emergence of MΦ^Prim^ progenitors, and later to their defective differentiation. These defects were associated with a disrupted expression of gene sets related to embryonic patterning and neurodevelopment. Lyl-1-deficiency also induced a reduced production of mature MΦ/microglia in the early brain, as well as a transient reduction of the microglia pool at midgestation and in the newborn.

We thus identify Lyl-1 as a critical regulator of MΦ^Prim^ and microglia development, which disruption may impair resident-MΦ function during organogenesis.

**Key points:** 1- Yolk sac primitive macrophage progenitors and microglia/Border Associated macrophages express Lyl-1.

2- Lyl-1-deficiency impairs primitive macrophage and microglia development and leads to the up-regulation of gene sets related to embryo patterning and neuro-development.

## INTRODUCTION

Amongst the components of the transcription factor network that regulate hematopoietic cells features, *Tal-1, Lmo2, Runx1* and *Gata2* stand out as major regulators of hematopoietic progenitor development.^*1, 2*^ *Tal-1, Lmo2* and *Gata-2* belong to a transcriptional complex, which also includes the basic helix-loop-helix (bHLH) transcription factor (TF) lymphoblastic leukemia-derived sequence 1 (*Lyl-1*). Unlike its paralog *Tal-1*, which is mandatory for the specification of all hematopoietic progenitors^*3, 4*^, Lyl-1 roles during developmental hematopoiesis remains poorly characterized. We analyzed these functions at the onset of YS hematopoietic development using *Lyl-1*^*LacZ/LacZ*^ mutant mice.^*5*^

During ontogeny, hematopoietic progenitors are generated in three successive and overlapping waves.^*6, 7*^ The emergence of the Hematopoietic Stem Cells (HSC) that will maintain lifelong hematopoiesis in the adult occurs at mid-gestation in the third and definitive hematopoietic wave. HSC generated in the aorta region immediately migrate to the fetal liver (FL) where they mature and amplify before homing to the bone marrow before birth.^*8, 9*^ Prior to HSC generation, the production of blood cells relies on two hematopoietic waves provided by the YS. This HSC-independent hematopoiesis comprises first the primitive hematopoietic wave, with the transient production of progenitors with embryonic specific features: From Embryonic day (E) 7.00, the YS produces monopotent progenitors for erythrocytes, megakaryocytes and macrophages (MΦ)^*10, 11*^, along with bipotent Erythro-Megakaryocytic (EMk) progenitors^*12*^, in a *Myb-*independent pathway.^*13, 14*^ The second YS wave, called transient-definitive, provides for a limited duration progenitors (mostly erythro-myeloid) that seed the FL and produce a hematopoietic progeny that displays definitive/adult differentiation features. Erythro-myeloid cell production in this wave occurs in a *Myb*-dependent pathway, through the progressive differentiation of erythro-myeloid progenitors (EMP) in a pathway similar to the adult one.^*6*^ As primitive and transient definitive YS waves both produce cells from erythro-myeloid lineages, they are also termed respectively “early EMP” and “late EMP”.^*15, 16*^

Considering the MΦ lineage, fate-mapping approaches aimed at determining the embryonic origin of resident-MΦs indicated that most tissues harbor resident-MΦs of diverse origins (YS, FL and adult bone marrow)^*15, 17, 18*^, which complicates the characterization of wave-dependent functions of the various subsets. However, these fate-mapping analyses established that, contrary to others tissue, brain MΦs (microglia and Border Associated MΦ (BAM)) develop only from YS-derived MΦ-progenitors^*14, 16, 19, 20, 21*^, confirming a model we previously put forward.^*22*^ Due to the coexistence of two waves in the YS, the origin of microglia has been debated (reviewed in ^*6, 15, 18, 23*^). An origin of microglia from MΦ-progenitors from the primitive/”early EMP” wave was supported by microglia labelling following an early (E7.0-E7.5) CRE-mediated induction of Runx1^*19*^ and by the intact microglia pool in *Myb*-deficient mice.^*14, 21*^ The origin of microglia was also attributed to the primitive wave in zebrafish embryos, since in MΦ^Prim^ progenitors arise in this species from a location distinct from other hematopoietic progenitors.^*24, 25, 26*^ Finally, the normal microglia development in mice lacking Kit ligand, leading to an impaired EMP development and the depletion of resident-MΦs in the skin, lung and liver supports this model.^*27*^

We here show that, at the early stages of YS hematopoiesis, *Lyl-1* expression characterizes primitive MΦ-progenitors. Through RNA-seq. analyses, it appears that these primitive MΦ-progenitors harbor an immune-modulatory phenotype, while those produce at a later stage favor the inflammatory signaling which promotes the emergence of HSC in the third and definitive hematopoietic wave. ^*28*^

Our results also indicate that in the brain, Lyl-1 is expressed in the entire microglia/BAM cell population at the onset of brain colonization and appeared to regulate microglia/BAM development.

Altogether, these data point to Lyl-1 as a major regulator of early embryonic MΦ-progenitors development and advocate for further analyses to more precisely delineate Lyl-1 function during the development of resident-MΦs in homeostatic and pathological contexts.

## RESULTS AND DISCUSSION

### Lyl-1 expression marks MΦ^Prim^ progenitors from the early YS

*Lyl-1* being expressed in the YS from the onset of YS hematopoiesis^*29*^, we first explored its function by characterizing the clonogenic potential of WT, *Lyl-1*^*WT/LacZ*^ and *Lyl-1*^*LacZ/LacZ*^ YS. E8-YS were maintained in organ culture for 1 day (E8 OrgD1-YS), allowing only the development of primitive and transient-definitive progenitors.^*30, 31*^ Compared to WT, the production of MΦ colonies was increased in *Lyl-1*^*WT/LacZ*^ and *Lyl-1*^*LacZ/LacZ*^ OrgD1-YS. Otherwise, the clonogenic potential and colony distribution were unmodified ***(Figure 1A)***.

**Figure 1:**
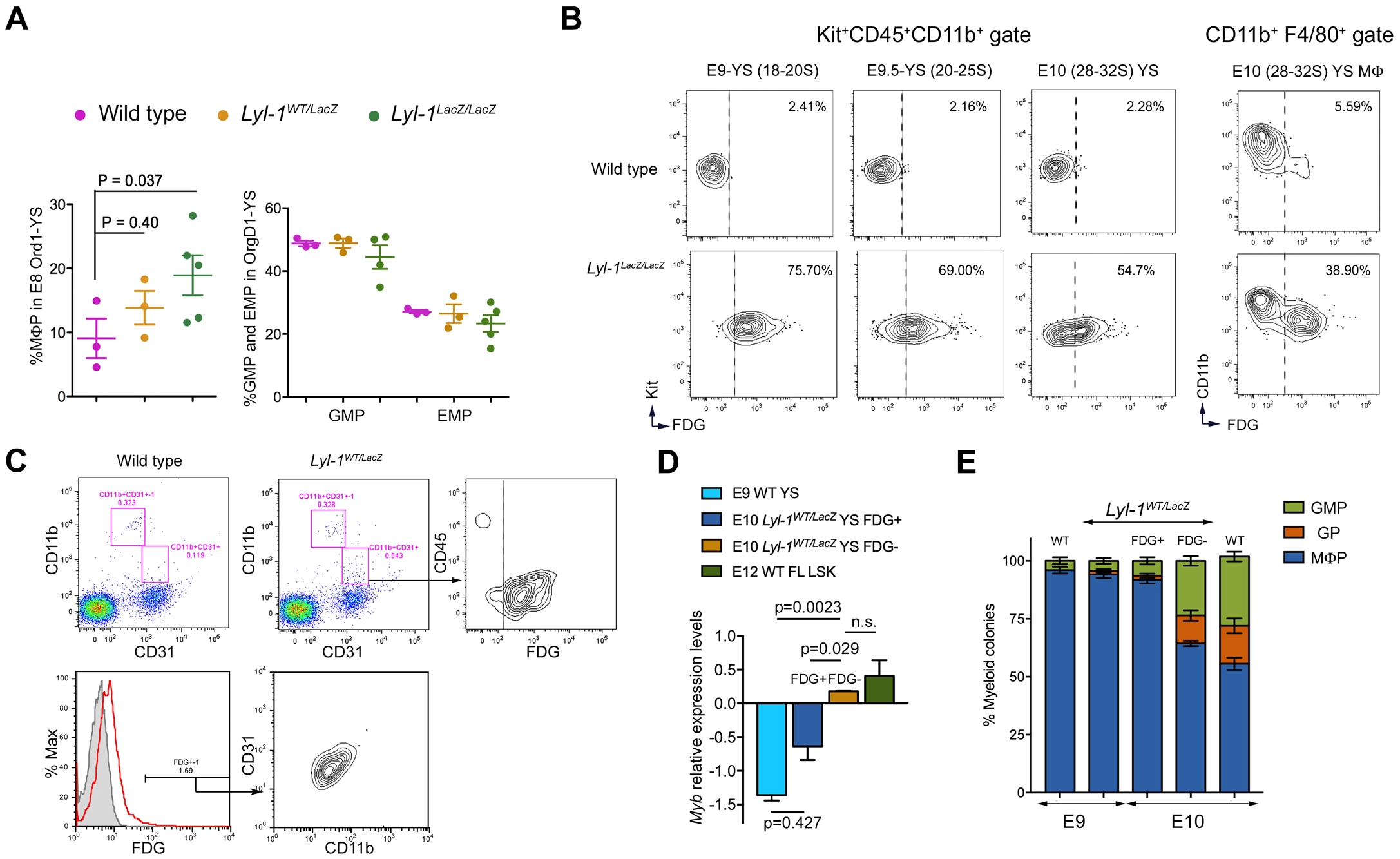
Lyl-1 expression marks MΦ^Prim^ progenitors in the early YS. **A. Lyl-1-deficiency leads to an increased production of MΦ-progenitors in the early YS:** Left: Clonogenic potential of E8 OrgD1-YS cells: production of MΦ-progenitors (CFU-M) in WT and *Lyl-1*^*LacZ/LacZ*^ OrgD1-YS (n=3-5, 3-6 YS per sample; mean ± s.e.m.; Unpaired, two-tailed *t*-Test). The size of the MΦ colonies and the cell morphology were similar for the 3 genotypes (data not shown). Right: distribution of other progenitors with a myeloid potential (EMP and GM) in WT, *Lyl-1*^*WT/LacZ*^ and *Lyl-1*^*LacZ/LacZ*^ E8 OrgD1-YS. **B. Lyl-1 expression in MΦ-progenitors:** FACS-Gal assay, using the β-Gal fluorescent substrate FDG was used as a reporter for Lyl-1 expression. While all MΦ-progenitors in E9-YS (left panel) expressed FDG/Lyl-1, E9.5 and E10-YS (middle panels) harbored two MΦ-progenitor subsets discriminated by their FDG/Lyl-1 expression. FDG^+^/Lyl-1^+^ and FDG^-^/Lyl-1^-^ mature MΦs (CD11b^+^F4/80^+^) also coexisted in E10-YS (right). The contour plots in WT samples indicate the level of non-specific background β-Gal activity/FDG labeling in WT samples. Representative profiles of 3 independent samples, each consisting of 3-4 YS *(See the gating strategy in* ***Supplemental figure 1A***). **C. MΦ**^**Prim**^ **progenitors express Lyl-1**. Upper panel: Flow cytometry profiles of WT (left) and *Lyl-1*^*WT/LacZ*^ (middle left) E8-YS (0-3S). CD11b^+^CD31^-^ MΦs (top gate) correspond to maternal MΦs present in E8-YS.^*11*^ All CD11b^+^CD31^+^ MΦ-progenitors (lower gate) displayed FDG/Lyl-1 expression. **D. RT-qPCR quantification of *Myb* expression levels**: Kit^+^CD45^+^CD11b^+^ progenitors were sorted from WT E9-YS, WT and *Lyl-1*^*WT/LacZ*^ E10-YS, and from FDG/Lyl-1 positive and negative fractions of MΦ-progenitors from *Lyl-1*^*WT/LacZ*^ E10-YS. Lin^-^Sca^+^Kit^+^ (LSK) progenitors from WT E12-FL were used as positive control. FDG^+^/Lyl-1^+^ MΦ-progenitors from E10-YS expressed *Myb*^*Low/Neg*^ levels similar to MΦ^Prim^ progenitors from E9-YS. The FDG^-^/Lyl-1^-^ fraction expressed significantly higher *Myb* levels, similar to LSK cells from E12-FL. *Myb* expression levels, shown on a Log^2^ scale, were normalized to the mean expression value obtained for WT E10-YS, considered as 1 (Unpaired, two-tailed *t*-Test). **E. FDG/Lyl-1 positive and negative myeloid progenitors produce a distinct progeny:** Clonogenic assays characterization of the type of progenitors produced by myeloid progenitors (Ter119^-^Kit^+^CD45^+^CD11b^+^) sorted from WT and *Lyl-1*^*WT/LacZ*^ E9-YS (<18 S; n=7) and E10-YS (n=15) in 3 independent experiments. At E10, myeloid progenitors from *Lyl-1*^*WT/LacZ*^ YS were subdivided into FDG/Lyl-1 negative (n=15) and positive (n=12) fractions (5 independent experiments). Samples were biological replicates comprising 6-8 YS. 100 to 150 Kit^+^CD45^+^CD11b^+^ cells per condition were platted in triplicate. All samples produced few non-myeloid contaminants, such as EMk and EMP in similar, non-significant amounts. FDG^+^/Lyl-1^+^ progenitors essentially produced MΦ colonies, while FDG^-^/Lyl-1^-^ progenitors produced also GM and G colonies.

Using FACS-Gal assay, we noticed that the entire MΦ-progenitor population (Kit^+^CD45^+^CD11b^+^) expressed Lyl-1 at E9. In contrast, two MΦ-progenitor subsets, discriminated by FDG/Lyl-1 expression, were present after E9.5 ***(Figure 1B)***. Since after E9.5, the YS harbors both MΦ^Prim^ and transient-definitive (MΦ^T-Def^) MΦ–progenitors, and as these progenitors subsets cannot be discriminated by phenotype,^*11*^ we investigated the known features discriminating these two waves: the origin from monopotent progenitors for MΦ^Prim^ progenitors^*10, 11*^ and the *Myb*-dependent^*14, 32*^ differentiation of MΦ^T-Def^ progenitors from EMP, via the production of granulo-monocytic (GM-), then granulocyte (G-) and MΦ-progenitors.^*6*^

At E8 (0-5 somites (S)), the YS only harbors MΦ^Prim^ progenitors, harboring a CD11b^+^CD31^+^ phenotype.^*11*^ At this stage, all MΦ-progenitors expressed FDG/Lyl-1 ***(Figure 1C)***. Most FDG^+^/Lyl-1^+^ CD11b and CD31cells (69.27%±0.33%) from *Lyl-1*^*WT/LacZ*^ E8-YS reliably produced MΦ colonies (72.78±9.65%; n=3) in clonogenic assays, amounting 1-4 MΦ progenitors per YS, a value consistent with previously published data.^*10, 11*^

Lyl-1 expression by MΦ^Prim^ progenitors was strengthened by RT-qPCR comparison of *Myb* expression: Both E9-YS MΦ^Prim^ progenitors and FDG^+^/Lyl-1^+^ progenitors from E10-YS expressed low *Myb* levels strengthening their primitive status, while FDG^-^/Lyl-1^-^ progenitors from E10-YS progenitors displayed *Myb* levels similar to lineage-negative Sca1^+^cKit^+^ progenitors from E12-FL ***(Figure 1D)***.

The differentiation potential of FDG^+^/Lyl-1^+^ and FDG^-^/Lyl-1^-^ fractions of Ter119^-^Kit^+^CD45^+^CD11b^+^ myeloid progenitors isolated from E10-YS also pointed to Lyl-1 expression by MΦ^Prim^ progenitors ***(Figure 1E)***: Similar to WT E9-YS MΦ^Prim^ progenitors, E10 FDG^+^/Lyl-1^+^ progenitors appeared monopotent, as they nearly exclusively produced MΦ colonies. In contrast, E10 FDG^-^/Lyl-1^-^ myeloid progenitors produced GM, G and MΦ colonies, a feature typical of transient-definitive progenitors ^*6*^. Overall, these data together suggested that Lyl-1 may mark MΦ^Prim^ progenitors from the earliest wave.

### Distinct features of WT MΦ-progenitors at E9 and E10

The distinction between E9 and E10 MΦ-progenitors was confirmed in RNA-seq. analysis of CD45^+^CD11b^+^Kit^+^ MΦ-progenitors sorted at E9 (MΦ^Prim^ progenitors) and E10 (MΦ^Prim^ and MΦ^T-Def^ progenitors). Principal Component Analysis separated E9 and E10 MΦ-progenitors according to stage and genotypes ***(Figure 2A)***. E9 and E10 WT MΦ-progenitors differed by the expression of 726 genes, 176 were up-regulated at E9 and 550 at E10. Considering the coexistence of MΦ^Prim^ and MΦ^T-Def^ progenitors in E10-YS, differentially expressed genes (DEGs) found at E10 may reflect wave-specific differences or stage-dependent changes related to MΦ^Prim^ progenitor maturation.

**Figure 2:**
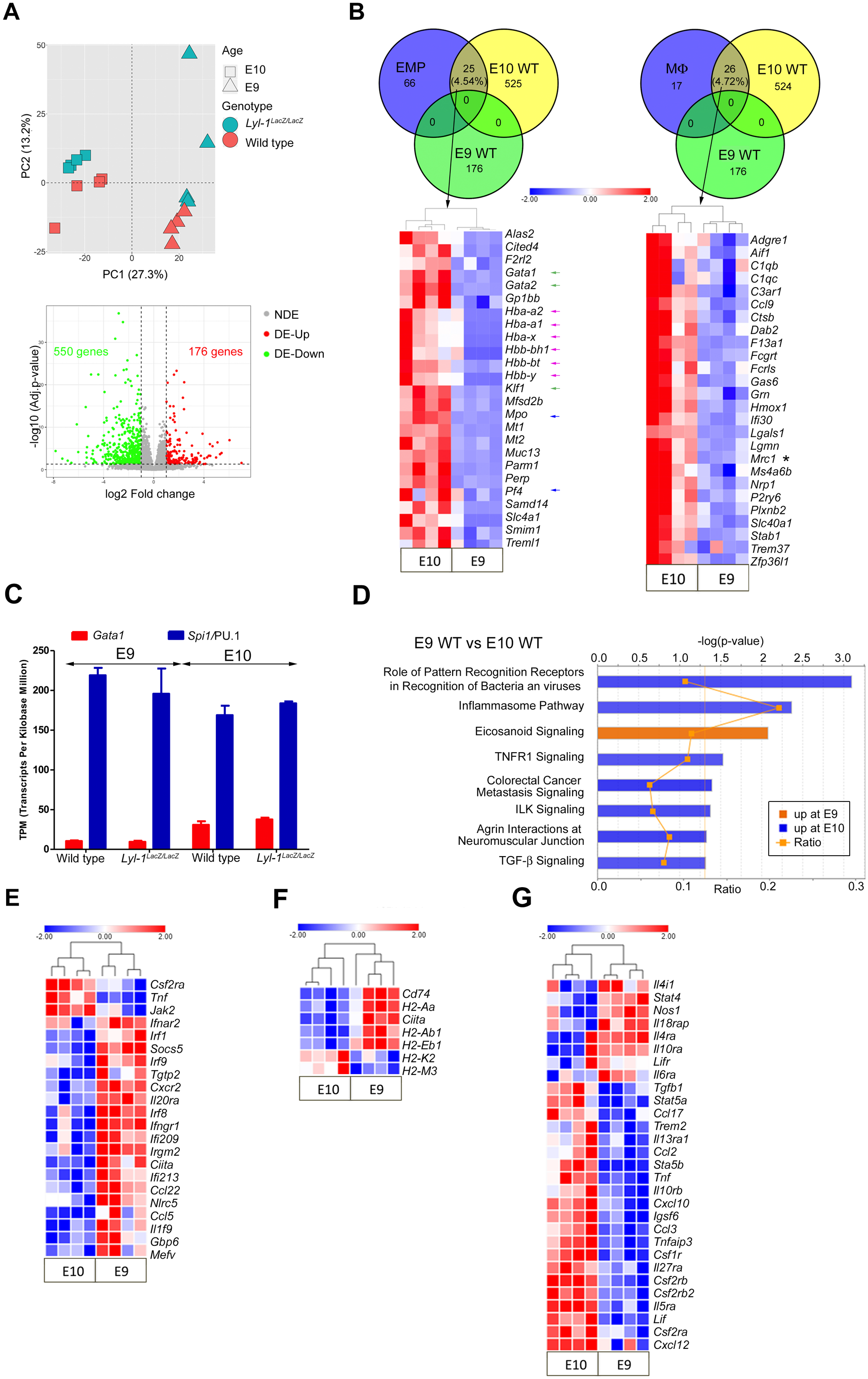
*Distinct features of WT* MΦ*-progenitors at E9 and E10*. **A**. Differentially expressed genes in MΦ-progenitors (Kit^+^CD45^+^CD11b^+^) sorted from WT and *Lyl-1*^*LacZ/LacZ*^ YS at E9 and E10. Upper panel: Unsupervised principal component analysis (PCA) plot positioned E9 and E10 MΦ-progenitors in two distinct groups, followed by segregation of WT and *Lyl-1*^*LacZ/LacZ*^ samples. Lower panel: Volcano plot of E9 WT vs E10 WT MΦ-progenitors. Red and green dots indicate genes with statistically significant changes in expression level. (*p*-value <0.05, absolute fold change≥2.0) (NDE: not deregulated genes; DE-Up: up-regulated genes; DE-Down: down-regulated genes). **B**. Upper panel: Venn diagram comparing DEGs in E9 WT versus E10 WT MΦ-progenitors to the EMP (Left) or MΦ signatures defined by Mass et al.^*33*^ (GEO accession number GSE81774). The number and percentage of DEGs common to the EMP or MΦ signatures is shown. Lower panel: Expression profiles of the overlapping genes identified in the Venn diagram (Heatmap displays transformed log2-expression values; Unpaired *t*-Test, two-tailed). Note the higher expression at E10 of genes involved in erythroid (Globins: Pink arrow; Transcription factors: green arrow), and megakaryocytic and granulocytic-related genes (blue arrow), and of *Mrc1*/CD206 (Asterisk). **C**. Relative expression levels of *Gata1* and *Spi1/*PU.1, indicated by their relative Transcripts per million kilo-bases (TPM). **D**. Enriched Pathways in E9 and E10 WT MΦ-progenitors with absolute z-score ≥2, from QIAGEN’s Ingenuity® Pathway Analysis (IPA). Bars: minus log of the p-value of each canonical pathway; Orange line: threshold p-value of 0.05. Ratio: genes detected/genes per pathway. **E**. Expression profiles of DEGs related to IFNγ and IFNβ response, identified by g:Profiler. (Heatmap displays transformed log2-expression values; unpaired t-Test, two-tailed). **F**. Expression profiles of DEGs related to MHC-II complex (Heatmap displays transformed log2-expression values; unpaired *t*-Test, two-tailed). **G**. Expression profiles of DEGs related to cytokine signaling (Heatmap displays transformed log2-expression values; unpaired *t-*Test, two-tailed).

Overlapping the identified DEGs to the EMP and E10.25-E10.5 MΦs signatures obtained by Mass *et al*.^*33*^ confirmed that WT E9 MΦ^Prim^ progenitors were distinct from these two populations, since none of the 176 upregulated at E9 belonged to these signatures. Comparatively, about 5% of the genes up-regulated at E10 belonged to the EMP and MΦ signatures ***(Figure 2B)***. A similar separation was observed in GSEA analyses ***(Supplemental figure 1B*)**. These observations suggest that within E10 MΦ-progenitors some, likely the MΦ^T-Def^ ones, retain part of the EMPs signature. In a WT context, E9 MΦ^Prim^ progenitors differed from E10 MΦ-progenitors by their TF repertoire. Genes regulating erythroid development (*Gata1, Gata2, Klf1*) and globin genes, embryonic (*Hbb-bh1, Hba-x, Hbb-y*) and definitive (*Hba-a2, Hba-a1, Hbb-bt*) were enriched at E10 ***(Figure 2B)***, while *Spi1*/PU.1 was highly expressed compared to *Gata1* at both stages ***(Figure 2C)***. The lower expression level at E9 of erythroid genes and of genes involved in granulo-monocytic (*Mpo, Csf2r/GM-CSF receptors, Cebp, Jun*) and megakaryocytic development (*Pf4, TPO signaling*) ***(Figure 2B; Supplemental table 1)*** sustains the monopotent/primitive status of E9 MΦ progenitors, and suggests that MΦ^T-Def^ progenitors may retain the expression of genes that characterize their EMP ancestor.

IPA and GSEA analyses indicated that E9 MΦ^Prim^ progenitors were more active in Eicosanoid signaling than E10 progenitors ***(Figure 2D)***. They were also enriched in type I interferon (IFN) β and type II IFNγ signaling ***(Figure 2E)*** and in MHC-II related genes, especially *Cd74* (top 1 IPA network) ***(Figure 2F; supplemental figure 1C)***. Cytometry analyses confirmed a low, but significant, enrichment of MHC-II expression at E9, compared to E10 ***(Supplemental figure 1D)***. Comparatively, E10 MΦ-progenitors were more active in inflammatory signaling ***(Figure 2D, E; supplemental figure 1C, E-G; supplemental table 1)***, and metabolically active ***(Supplemental table 1)***. The complement cascade and phagocytosis also prevailed at E10 ***(Supplemental figure 1H-I)***.

Altogether, the signature for WT E9 MΦ^Prim^ progenitors points to an immuno-modulatory function, while E10 MΦ-progenitors appear involved in phagocytosis and inflammatory signaling. Interestingly, inflammatory signaling has been revealed as a key factor favoring embryonic HSC emergence (reviewed in^*34*^). The source of inflammatory signals was further identified as MΦ progenitors expressing *Mrc1*/CD206^*35*^, a marker up-regulated in E10 WT MΦ-progenitors compared to E9 ***(Figure 2B)***.

### Lyl-1 deficiency impacts embryonic development

When evaluating the effect of *Lyl-1*-deficiency at the earliest stage of MΦ^Prim^ development, clonogenic assays pointed to an increased production of MΦ-progenitors in Mutant E8-YS compared to WT ***(Figure 3A)***, concordant with our first observation ***(Figure 1A)***. At this stage, the increased size of the initial MΦ-progenitor pool appeared to results from an elevated commitment of mesodermal/pre-hematopoietic cells to a MΦ fate, rather than from a defective differentiation ***(Supplemental figure 2> A-D and related information)***. The high increase of *Itga2b*/CD41 expression level in E9 *Lyl-1*^*LacZ/LacZ*^ MΦ^Prim^ progenitors ***(Figure 3B)*** may reflect this elevated commitment. Lyl-1 expression in YS mesoderm^*29*^, where it cannot substitute for *Tal-1* mandatory function for the generation of hematopoietic progenitors^*4*^, was already established^*3*^. Recently, Lyl-1 was identified as a regulator of mesoderm cell fate^*36*^ and of the maintenance of primitive erythroid progenitors.^*37*^

**Figure 3:**
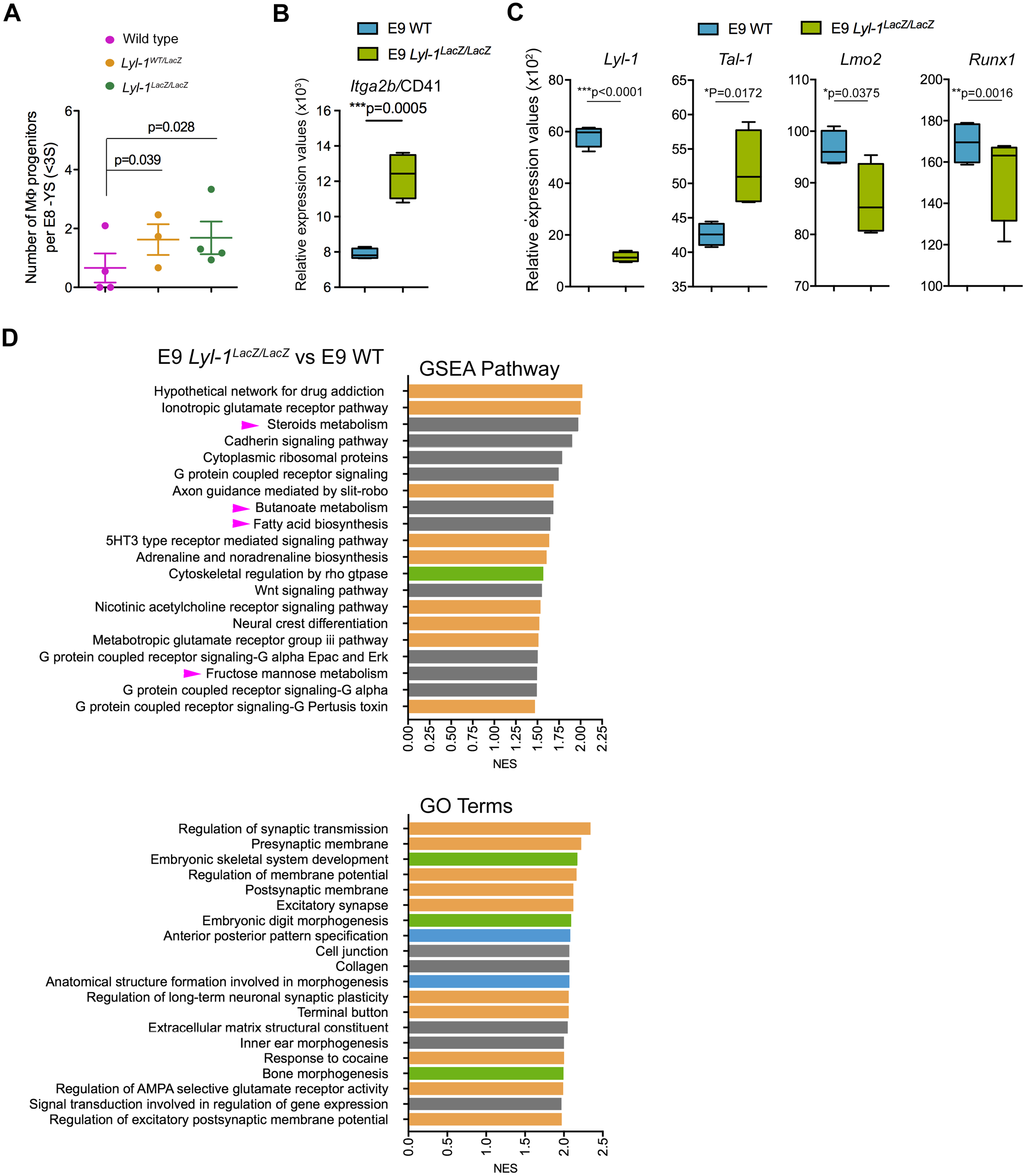
Lyl-1 regulates the production of E8 MΦ^Prim^ progenitors. **A**. In clonogenic assays, the number of MΦ colonies obtained from E8-YS (0-3S) was increased in *Lyl-1*^*WT/LacZ*^ and *Lyl-1*^*LacZ/LacZ*^ compared to WT. The majority of the 25-30 colonies per YS were Ery^P^ (60 to 80% in each 3 genotypes). Other progenitors were occasionally and randomly detected in WT and mutant samples (less than one EMP (0.81%±0.66; n=3) and/or GM progenitor per E8-YS), confirming that the assay was performed at a time when MΦ^T-Def^ progenitors were absent. (n=3-5, 5-10 YS per sample; plots show mean ± s.e.m.; Unpaired, two-tailed *t*-Test). **B**. Relative expression levels of the CD41 coding gene *Itg2b* in WT and *Lyl-1*^*LacZ/LacZ*^ MΦ-progenitors at E9 (unpaired *t*-Test, two-tailed). **C**. Relative expression levels of TF regulating hematopoietic progenitor emergence in *Lyl-1*^*LacZ/LacZ*^ MΦ-progenitors compared to WT at E9 (unpaired, two-tailed *t*-Test). **D**. GSEA pathways (Top; FDR q-Value <0.29) and GO terms (Bottom; FDR q-value <0.01) enriched in E9 *Lyl-1*^*LacZ/LacZ*^ compared to E9 WT MΦ-progenitors. Highlighted are the pathways specifically related to embryo patterning (blue) and to the development of skeletal (green) and nervous systems (yellow). Pink arrows point to changes related to metabolic pathways.

The TF network that controls developmental hematopoiesis^*2*^ was also modified ***(Figure 3C)***: beside the expected reduction of *Lyl-1* expression, the expression of *Lmo2*, a *Lyl-1* target^*38*^, was down-regulated, while *Tal-1* up-regulation might reflect some compensatory function.^*3*^ The consequences were apparent in GSEA analyses: both pathways and GO terms uncovered an up-regulation of signaling pathways involved in embryo patterning (Wnt, Hox and Smad) in *Lyl-1*^*LacZ/LacZ*^ MΦ-progenitors, as well as a highly modified collagen, integrin and cadherin usage ***(Supplemental table 2)***. Accordingly, developmental trajectories were affected ***(Figure 3D)***, with the up-regulation in E9 *Lyl-1*^*LacZ/LacZ*^ MΦ-progenitors of gene sets related to “anterior-posteriorpattern specification” and “anatomical structure formation involved in morphogenesis”, notably skeletal and nervous system development.

GSEA and KEGG comparison of *Lyl-1*^*LacZ/LacZ*^ and WT MΦ-progenitors at E10 highlighted another patterning modification, namely the down-regulation of gene sets involved in heart development ***(Supplemental figure 2E; supplemental table 3)***, which might stem from a defective MΦ development. The heart harbors three resident-MΦ subsets, two of which originate from the YS.^*39*^ Amongst the features that distinguish WT E9 MΦ^Prim^ progenitors from E10 MΦ-progenitors, the enriched expression of MHC-II ***(Figure 2F)*** and poor expression of phagocytosis-related genes ***(Supplemental figure 1I)*** at E9 also characterize one of the two YS-derived CCR2^-^ resident-MΦ subsets in the heart.^*39, 40*^ Therefore, a function for Lyl-1^+^ MΦ^Prim^ progenitors in heart development may be considered. This observation reinforces the need to better characterize the contributions of MΦ-progenitors from both primitive and transient-definitive waves to tissues harboring YS-derived resident-MΦs.

The patterning defects highlighted in defective MΦ^Prim^, might be responsible, at least in part, for the significantly decreased litter size and increased perinatal lethality observed in *Lyl-1*^*LacZ/LacZ*^ mice compared to WT ***(Supplemental figure 2F)***, which indicates the requirement for Lyl-1 during various developmental processes.

### Defective MΦ^Prim^ development in Lyl-1 mutant YS

The analysis of *Lyl-1* expression in A1-A2-A3 subsets from *Cx3cr1*^*WT/GFP*^ YS indicated that *Lyl-1* is expressed throughout MΦ-progenitor differentiation, with levels decreasing from A1 to A3 ***(Supplemental figure 3A)***. We monitored the distribution of A1-A2 and A3 MΦ subsets ***(Supplemental figure 2A)*** at E10-YS, when all three subsets are present, using the *Cx3cr1*^*WT/GFP*^*:Lyl-1*^*LacZ/LacZ*^ strain. While the size of the whole MΦ population was not overtly modified, *Lyl-1*-deficiency impacted the subset distribution, with increased A1 and reduced A2 and A3 pool sizes ***(Figure 4A)***. Lyl-1 appears to regulate MΦ-progenitor differentiation towards mature MΦs. This defect could result from the altered cytokine signaling uncovered in E9 mutant progenitors through GSEA and IPA analyses ***(Figure 4B; supplemental figure 3B)***. A limited or delayed differentiation of E9 MΦ^Prim^ progenitors was supported by the down-regulated *Spi1*/PU.1 signaling pathway ***(Supplemental figure 3C; Supplemental table 4B)*** and the decreased expression of *Ptprc*/CD45, *Csfr1, Itgam*/CD11b and CD33 ***(Figure 4C)***.

**Figure 4:**
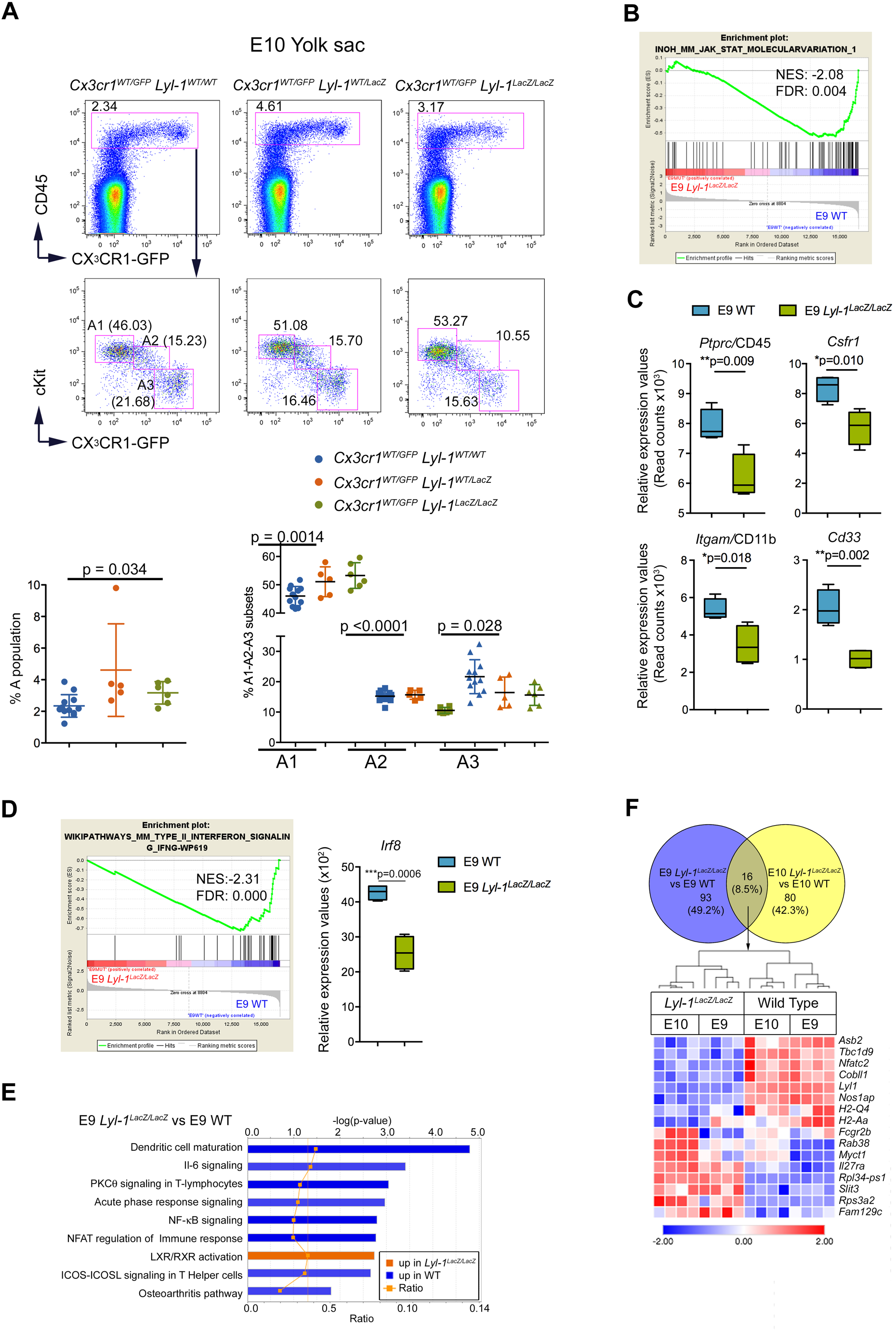
Defective differentiation of Lyl-1^LacZ/LacZ^ MΦ-progenitors from E10-YS. **A**. Distribution of A1-A2 and A3 MΦ subsets in E10-YS from *Cx3cr1*^*WT/GFP*^*:Lyl-1*^*WT/WT*^, *Cx3cr1*^*WT/GFP*^*:Lyl-1*^*WT/LacZ*^ and *Cx3cr1*^*WT/GFP*^: *Lyl-1*^*LacZ/LacZ*^ embryos. While the size of the whole MΦ population is similar in the three genotypes (Top panel), *Lyl-1* deficiency leads to a modified distribution of the MΦ subsets (middle and lower panel) with an increased size of the A1 subset and a reduced A3 pool (5-12 independent analyses, 6-8 YS per sample. Plots show mean ± s.e.m.; Unpaired, two-tailed *t*-Test). **B**. GSEA pathway indicates a deficit in Jak1-Stat signaling in *Lyl-1*^*LacZ/LacZ*^ MΦ-progenitors compared to WT at E9 (NES: normalized enrichment score; FDR: false discovery rate). **C**. Relative expression levels (read counts) of hematopoietic markers in WT and *Lyl-1*^*LacZ/LacZ*^ MΦ-progenitors at E9 (unpaired *t*-Test, two-tailed). **D**. Top 1 GSEA pathway indicates that the IFN signaling pathway (left) which characterize E9 MΦ^Prim^ progenitors, and particularly *Irf8* (right), is defective in *Lyl-1*^*LacZ/LacZ*^ MΦ-progenitors (NES: normalized enrichment score; FDR: false discovery rate). **E**. From the 53 canonical pathways identified by IPA in the DEGs, 9 were enriched with an absolute Z score ≥ 1. Bars: minus log of the *p*-value of each canonical pathway; Orange line: threshold p-value of 0.05. Ratio: genes detected/genes per pathway. **F**. Upper panel: Venn diagram comparing the DEGs in E9 *Lyl-1*^*LacZ/LacZ*^ vs E9 WT to those in E10 *Lyl-1*^*LacZ/LacZ*^ vs E10 WT MΦ-progenitors. Lower panel: Expression profiles of the DEGs common to both stages identified by the Venn comparison (Heatmap displays transformed log2-expression values; unpaired *t*-Test, two-tailed).

Within the MΦ lineage, Lyl-1 function during normal development would initially consist to restrict the size of the MΦ^Prim^ progenitor pool and/or the duration of its production, which is transient^*6*^, as indicated by the maintenance of the intermediate mesoderm to MΦ-progenitor pool observed in *Lyl-1*^*LacZ/LacZ*^ E8-YS. Indeed, the increased size of the MΦ-progenitor pool in E8-E9 YS appears independent from the defective/delayed differentiation of MΦ-progenitors observed at E10, since this process starts after E9.5.^*10, 11, 15*^ Subsequently, the increased size of MΦ^Prim^ progenitor pool in E10 *Lyl-1*^*LacZ/LacZ*^ YS likely results from a defective/delayed differentiation mediated by a defective cytokine signaling, implying that during normal development, Lyl-1 would promote the differentiation of MΦ^Prim^ progenitors.

*Lyl-1*^*LacZ/LacZ*^ MΦ-progenitors were also deficient in the IFN signaling that characterize E9 MΦ^Prim^ progenitors, notably *Irf8*, a factor involved in YS-MΦ and microglia development^*21, 41*^ ***(Figure 4D)***. Compared to WT, MΦ-progenitors from *Lyl-1*^*lacZ/lacZ*^ E9-YS up-regulated the LXR/RXR activation pathway ***(Figure 4E)*** and metabolic pathways, some enriched WT MΦ-progenitors at E10 (Butanoate, steroid) ***(Supplemental table 1)***, and other not (Fructose/mannose, fatty acid) ***(Figure 3D)***. They were also less active in inflammatory signaling pathways, particularly through NFκb, a factor known to interact with *Lyl-1*^*42*^, and in TLR signaling ***(Figure 4D; supplemental figure 3B, D-E; supplemental table 4B)***. Overlapping the DEGs identified in *Lyl-1*^*LacZ/LacZ*^ MΦ-progenitors at E9 and E10 identified the core signature of *Lyl-1*-deficiency, independent of the maturation occurring between these stages ***(Figure 4F)***.

Unfortunately, the co-existence of MΦ^Prim^ and MΦ^T-Def^ progenitors in E10-YS complicates the attribution of gene expression changes to a stage-dependent maturation of MΦ^Prim^ progenitors or to a signature specific to MΦ^T-Def^ progenitors. However, most pathways favored by E10 progenitors were insensitive to *Lyl-1*-deficiency, except TLR signaling pathway that was down-regulated in E9 *Lyl-1*^*LacZ/LacZ*^ progenitors, compared to WT.

### Lyl-1-expressing MΦ-progenitors contribute to the fetal liver and brain

The FL^*9*^ and brain^*19, 22*^ are colonized as early as E9 by YS-derived resident-MΦ progenitors. We evaluated the contribution of Lyl-1-expressing MΦ^Prim^ progenitors to these rudiments at E10 ***(Figure 5A)***. While E10-YS comprised FDG^+^/Lyl-1^+^ and FDG^-^/Lyl-1^-^ MΦ-progenitors and mature (F4/80^+^) MΦ subsets ***(Figure 1B)***, the brain from the same embryos essentially harbored FDG^+^/Lyl-1^+^ MΦ-progenitors and MΦs. In contrast, both FDG^+^/Lyl-1^+^ and FDG^-^/Lyl-1^-^ MΦ-progenitors were present in E10-FL, as in E10-YS, and MΦ-progenitors were more abundant in mutant FL than in WT ***(Figure 5B)***.

**Figure 5:**
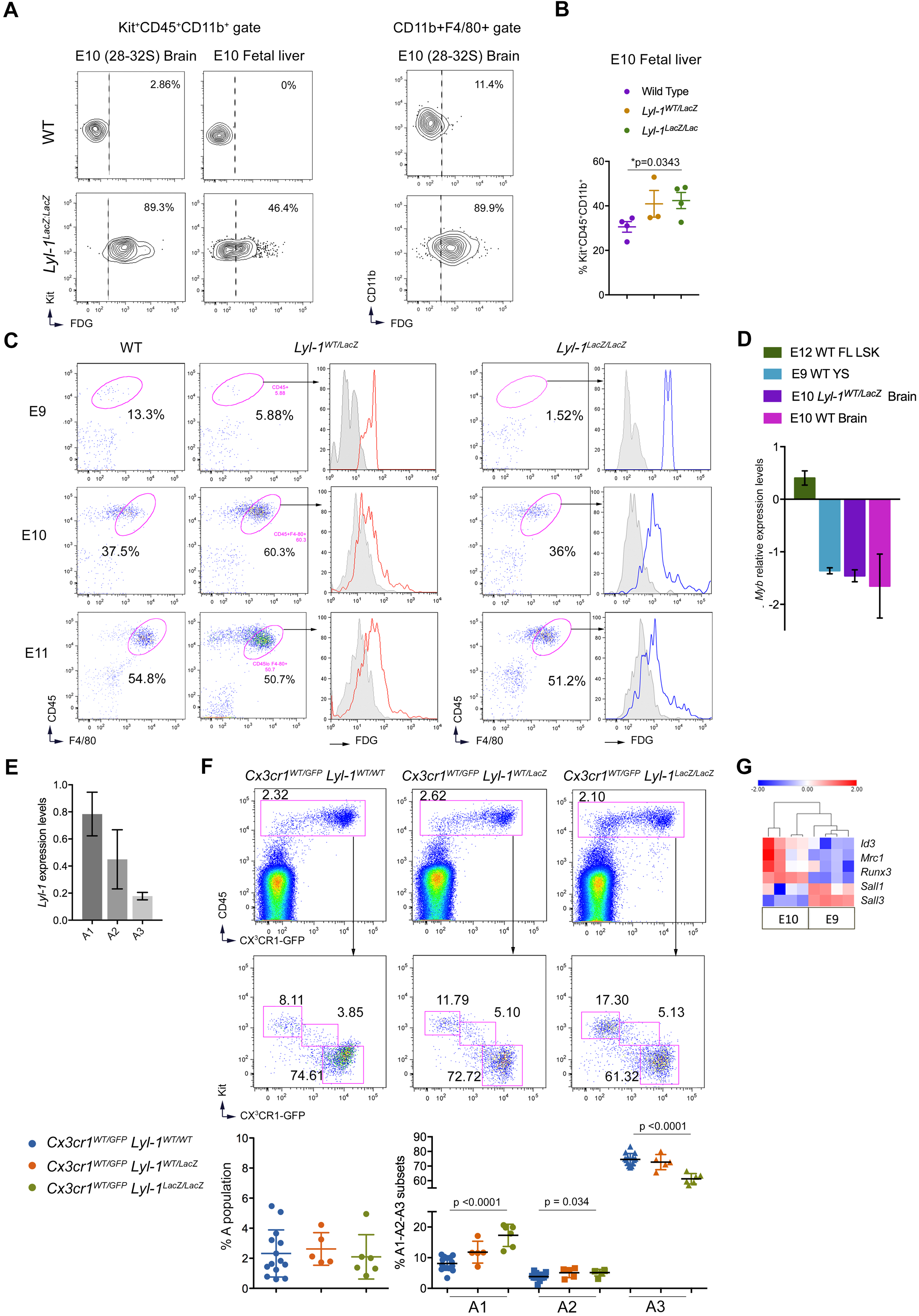
Contribution of Lyl-1-expressing MΦ-progenitors to the fetal liver and brain. **A**. Left panel: All MΦ-progenitors from E10-Brain (Left) expressed Lyl-1, contrary to the corresponding YS *(Figure 1B right)* which harbor both FDG^+^/Lyl-1^+^ and FDG^-^/Lyl-1^-^ subsets. MΦ-progenitors from E10 *Lyl-1*^*LacZ/LacZ*^ FL (right) harbored both FDG^+^/Lyl-1^+^ and FDG^-^/Lyl-1 MΦ-progenitor subsets. Right panel: in E10-brain, mature MΦs (CD11b^+^F4/80^+^ gate) were all FDG^+^/Lyl-1^+^. The contour plots in WT samples tTop panel) indicate the level of non-specific background β-Gal activity/FDG labeling in WT samples. Representative profiles of 3 independent samples, each consisting of 3-4 E10-Brain or 8-12 E10-FL. **B**. Quantification of cKit^+^CD45^+^CD11b^+^ MΦ-progenitors in E10-FL (plots show mean ± s.e.m.; Unpaired, two-tailed *t*-Test). **C. Lyl-1 marks the entire F4/80+ microglia/BAM population from the onset of brain colonization**. FDG/Lyl-1 expression in F4/80+ microglia/BAM from the brain of *Lyl-1*^*WT/LacZ*^ and *Lyl-1*^*LacZ/LacZ*^ embryos at E9 to E11. The rare CD11b^+^ F4/80^low-neg^ cells present in the brain at E9 are FDG^+^/Lyl-1^+^ (Top Panel). Grey histograms indicate non-specific background β-Gal activity/FDG levels in WT samples. **D. MΦ-progenitors from E10-brain express *Myb* levels similar to E9-YS MΦ**^**Prim**^ **progenitors**. RT-qPCR quantification of *Myb* expression levels in Kit^+^CD45^+^CD11b^+^ MΦ-progenitors sorted from WT E9-YS and from WT and *Lyl-1*^*WT/LacZ*^ brain at E10. Lin^-^Sca^+^cKit^+^ (LSK) progenitors from WT E12-FL were used as positive control. *Myb* expression levels, shown on a Log^2^ scale, were normalized to the mean expression value obtained for WT E10-YS, considered as 1 (Unpaired, two-tailed *t*-Test). **E**. Heatmap expression profile of the genes that mark the development of tissue resident-MΦs in WT E9 and E10 MΦ-progenitors (Heatmap displays transformed log2-expression values; Unpaired, two-tailed *t*-Test). **F**. RT-qPCR analyses of *Lyl-1* expression in A1 to A3 MΦ subsets isolated from *Cx3cr1*^*WT/GFP*^ brain at E10. *Lyl-1* is expressed by the 3 subsets, with levels decreasing with differentiation. Expression levels were normalized to the mean value obtained for *Cx3cr1*^*WT/GFP*^ YS A1 progenitors (n=3). **G. Defective differentiation of brain MΦ-progenitor in Lyl-1 mutant embryos**. Distribution of A1-A2 and A3 MΦ subsets in E10 brain from *Cx3cr1*^*WT/GFP*^*:Lyl-1*^*WT/WT*^, *Cx3cr1*^*WT/GFP*^*:Lyl-1*^*WT/LacZ*^ and *Cx3cr1*^*WT/GFP*^*:Lyl-1*^*LacZ/LacZ*^ embryos. The size of the whole MΦ population was similar in the three genotypes (Top panel), but Lyl-1 deficiency modified the distribution of the MΦ subsets (middle and lower panel) with an increased size of the A1 subset and a reduced A3 pool (5-12 independent analyses, 6-8 brains per sample. Plots show mean ± s.e.m.; Unpaired, two-tailed t Test).

We next focused on brain MΦs during the colonization stage, which lasts until E11.^*43*^ At this stage, microglia and perivascular, meningeal and choroid plexus MΦσ, collectively referred to as BAMs, are all located in the brain mesenchyme and therefore undistinguishable.^*16, 44*^ FACS-Gal assay demonstrated that the whole F4/80^+^ microglia/BAM expressed Lyl-1 throughout the settlement period ***(Figure 5C)***. The presence of FDG^+^/Lyl-1^+^F4/80^+^ microglia/BAM at early stage of brain colonization suggests that MΦs could participate to this step.

Lyl-1^+^ MΦ^Prim^ progenitors and early microglia/BAM shared similar features, such as an early appearance timing and low level of *Myb* expression ***(Figure 5D)***, concordant with a *Myb*-independent development of microglia.^*14, 21*^ *Lyl-1* was also similarly expressed in A1-A2 and A3 MΦ subsets from the YS and brain ***(Figure 5E; supplemental figure 3A)***. *Lyl-1*-deficiency impacted the distribution of MΦs subsets in E10 *Cx3cr1*^*WT/GFP*^*:Lyl-1*^*LacZ/LacZ*^ brain: an increased A1 and a reduced A3 pool size indicated that Lyl-1 regulates MΦ-progenitor differentiation in both YS and brain ***(Figure 5F)***.

The proximity between YS MΦ^Prim^ progenitors and microglia was also apparent in RNA-seq. data: E9 WT MΦ^Prim^ progenitors expressed significantly lower *Mrc1*/CD206 and higher *Sall3* levels than E10 MΦ-progenitors, and a slightly increased *Sall1* level ***(Figure 5G)***, a transcriptomic pattern that characterizes microglia.^*33, 43, 45*^ This partial bias toward a microglia signature suggests that the first stage of microglia development program is already initiated in MΦ^Prim^ /”early EMP”progenitors in E9-YS.

### Lyl-1 inactivation impairs microglia development at two development stages

Having defined Lyl-1 implication during microglia/BAM settlement in the brain, we turned to later development stages. Cytometry and database analyses^*43*^ confirmed the continuous expression of Lyl-1 in CD45^low^ microglia until adulthood ***(Supplementary figure 4A)***. *LYL-1* expression was also reported in microglia from healthy murine and human adults.^*46, 47, 48, 49*^ We examined the impact of *Lyl-1*-deficiency on microglia pool size during development. Microglia quantification pointed to E12 as the first step impacted. The arrested increase of microglia pool in *Lyl-1*^*LacZ/LacZ*^ brain at E12 ***(Figure 6A)*** resulted from a reduced proliferation ***(Figure 6B)*** rather than an increased apoptosis ***(Supplementary figure 4B)***. Moreover, *Lyl-1*-deficiency provoked morphological changes in E12 *Cx3cr1*^*WT/GFP*^*:Lyl-1*^*LacZ/LacZ*^, compared to *Cx3cr1*^*WT/GFP*^ microglia, namely a reduced number and extent of ramifications ***(Figure 6C; supplementary figure 4C-D)***. From E14, the microglia pool size returned to levels similar to WT ***(Figure 6A)***, probably due to the highly reduced apoptosis level in *Lyl-1*^*LacZ/LacZ*^ microglia at E14 ***(Supplementary figure 4B)***.

**Figure 6:**
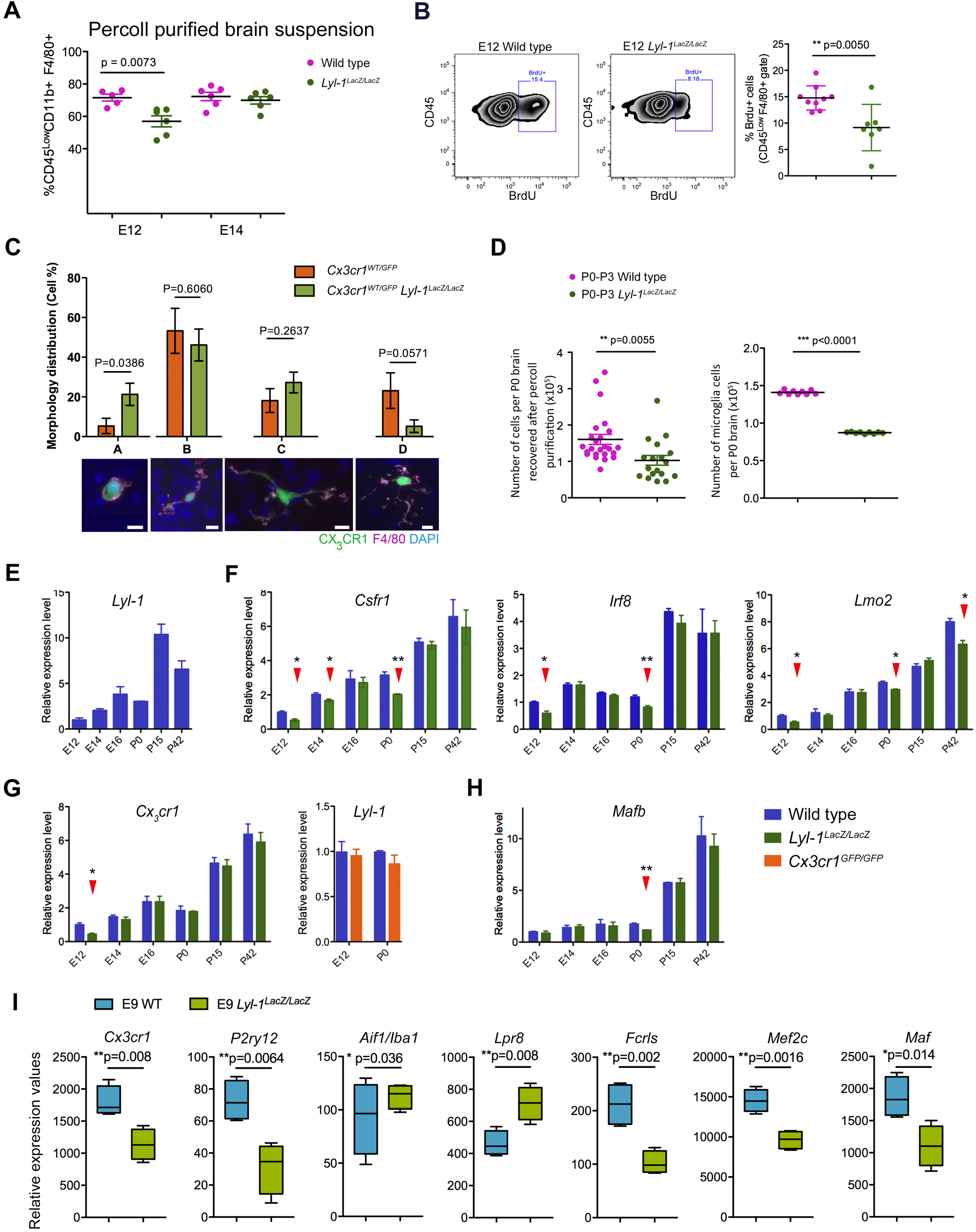
Lyl-1 deficiency leads to transient reductions of the microglia pool at E12 and P0-P3. **A**. Quantification of the microglia population in E12 and E14 brain showing the decreased size of the microglia pool at E12 and its recovery to a normal pool size at E14. Plots show mean ± s.e.m.; Two tailed, unpaired *t*-test. **B**. The decreased microglia pool at E12 may result from a reduced proliferation, as shown by the two folds decrease (right) of BrdU-labeled cells in *Lyl-1*^*LacZ/LacZ*^ (middle) compared to WT (left) brains. Plots show mean±s.e.m.; Two tailed, unpaired *t*-test. **C**. At E12, *Cx3cr1*^*WT/GFP*^*:Lyl-1*^*LacZ/LacZ*^ microglia displayed a reduced number and extent of ramifications compared to their *Cx3cr1*^*WT/GFP*^ counterpart. Bottom: Microglia morphology was classified into subtypes depending on the number of main ramifications (A: none, B: 2, C: 3 and D:>3). Top: Microglia deprived of ramifications predominated in *Lyl-1*-deficient microglia. 65 and 61 cells were respectively acquired from the midbrain of E12 *Cx3cr1*^*WT/GFP*^ and *Cx3cr1*^*WT/GFP*^*:Lyl-1*^*LacZ/LacZ*^ embryos (for each genotype, brains from 12 embryos were acquired in 3 independent experiments). Microglia were identified by *Cx3cr1*-driven GFP expression and F4/80-APC immuno-staining. Bar=10μm. Plots show mean±s.e.m.; Two tailed, unpaired *t*-test. **D**. In *Lyl-1*^*LacZ/LacZ*^ newborns, the cellularity of the brain was consistently lower than in WT (left), and so was the estimated microglia number (right). Plots show mean ± s.e.m.; Two tailed, unpaired *t*-test. **E**. Kinetic evolution of *Lyl-1* expression levels in WT microglia from embryonic stages to adulthood. An increased expression of *Lyl-1* from embryonic stages to adulthood was also inferred from timeline RNA-seq. data.^*43*^ (GEO accession number GSE79812). **F**. Quantitative RT-PCR analyses also point to E12 and P0 as key development stages regulated by Lyl-1. CD11b^+^F4/80^+^CD45^low^ microglia were isolated at sequential development stages. Bar graphs show the kinetic of expression of genes modified in *Lyl-1*^*LacZ/LacZ*^ microglia (arrowheads), normalized to the mean expression value in WT E12 microglia (n=3). Error bars indicate s.e.m. Two tailed, unpaired *t*-test. **G**. *Cx3Cr1* and *Lyl-1* expression in mutant microglia. The expression level of *Cx3CR1*, analyzed as in F, was decreased in Lyl-1 mutants at E12 (left), while *Lyl-1* expression levels were unmodified in *CX3CR1*^*GFP/GFP*^ microglia at E12 and in newborns (right). **H**. *Mafb* expression in mutant microglia. *Mafb* expression level, analyzed as in F, was reduced in the microglia of *Lyl-1*^*LacZ/LacZ*^ newborns. **I**. The expression of genes enriched in microglia and/or essential for their function are deregulated in *Lyl-1*^*LacZ/LacZ*^ MΦ-progenitors at E9. Relative expression levels (read counts) in WT and *Lyl-1*^*lacZ/lacZ*^ MΦ-progenitors from E9-YS (Unpaired, two-tailed *t*-Test).

P0-P3 was identified as a second stage altered in *Lyl-1*^*LacZ/LacZ*^ microglia. At birth, the cellularity of *Lyl-1*^*LacZ/LacZ*^ brain was significantly decreased compared to WT ***(Figure 6D)***, which was not the case at earlier stages ***(Supplementary figure 4E)***. CD11b^+^ cells recovery was also reduced (WT: 140.96±0.91×10^3^, n=9; *Lyl-1*^*LacZ/LacZ*^: 87.18±0.37×10^3^, n=9). Consequently, *Lyl-1*-deficiency triggered a nearly 2-fold reduction of the microglia population ***(Figure 6D)***. This perinatal reduction of microglia appeared transient, since no difference with WT brain was observed in the adult ***(Supplementary figure 4F)***. Transient decreases of microglia pool size, such as those we observed at E12 and P0-P3 in *Lyl-1*^*LacZ/LacZ*^ mutant, have been reported to occur during normal development in postnatal weeks 2-3^*50*^, but also in *Cx3cr1* mutant mice during the 1^st^ postnatal week^*51*^. This indicates a highly dynamic control of the microglia pool size during key steps of neural development that seems preserved in *Lyl-1* mutant, with the exception of the E12 and P0-P3 time-points. At this later stage, the reduction of brain cellularity in *Lyl-1*^*LacZ/LacZ*^ mice points to Lyl-1 as a possible regulator of the trophic function of microglia on brain cells^*52, 53*^.

The identification of E12 and P0-P3 as key stages for Lyl-1 function in microglia development was confirmed by RT-qPCR analyses of the expression of genes essential for MΦs (*Spi1/*PU.1, *Csf1r, Mafb*) and/or microglia (*Runx1, Cx3cr1, Irf8*) development and function, of known regulators of developmental hematopoiesis (*Tal-1, Lmo2, Runx1*) and related factors (*Tcf3*/E2A, *Tcf4*/E2.2) ***(Figure 6E-F; supplementary figure 4G)***. Time-course analyses highlighted the down-regulation of *Csf1r, Irf8* and *Lmo2* in *Lyl-1*^*LacZ/LacZ*^ microglia at E12 and P0-P3, while *Cx3cr1* was only decreased at E12 ***(Figure 6G)***. Note that *Lyl-1* expression was unmodified in *Cx3cr1*^*GFP/GFP*^ mutants ***(Figure 6G)***.Interestingly, *Cx3cr1*, as well as *Irf8* and *Lmo2*, belong to potential Lyl-1 target genes.^*2*^ *Mafb* expression levels in *Lyl-1*^*LacZ/LacZ*^ microglia transiently decreased at P0-P3 and later returned back to WT expression levels ***(Figure 6H)***. As Mafb represses resident-MΦ self-renewal^*54*^, the recovery of a normal amount of microglia after birth may stem from this transient decrease. *Spi1/*PU.1, *Tcf3*/E2A and *Tcf4*/E2.2 expression levels were unmodified in *Lyl-1*^*LacZ/LacZ*^ microglia, while *Runx1* expression was only affected after birth. *Tal-1* expression was decreased at E14 and increased after birth, suggesting that this *Lyl-1* paralog^*3*^ does not compensate *Lyl-1*-deficiency during embryonic stages, but may do so at postnatal stages ***(Supplementary figure 4G)***. Remarkably, RNA-seq. results indicated that some genes deregulated in *Lyl-1*^*LacZ/LacZ*^ microglia at E12 and P0-P3 were also down-regulated in *Lyl-1*^*LacZ/LacZ*^ MΦ-progenitors at E9 (*Csfr1:* ***Supplementary figure 3C***, *Lmo2:* ***Figure 3C***; *Irf8:* ***Figure 4D***; *Cx3cr1:* ***Figure 6I)***. These deregulations were transient, however in both locations and stages they coincided with a defective MΦ/microglia differentiation.

Other genes enriched in microglia (*Fcrls, Mef2c, Maf*)^*55*^ or involved in the maintenance of microglia homeostasis (*P2ry12*)^*56*^ were also expressed in E9 *Lyl-1*^*LacZ/LacZ*^ MΦ-progenitors at a lower level than in the WT, except *Lpr8* and *Aif-1/*Iba1 ***(Figure 6I)***. These deregulations highlight again shared features between MΦ^Prim^ progenitors and microglia/neural development which became apparent upon *Lyl-1* inactivation considering the large number of neural signaling pathways up-regulated in E9 *Lyl-1*^*LacZ/LacZ*^ MΦ-progenitors ***(Figure 3D)*** and the relationship of the DEGs enriched in E9 MΦ^Prim^ progenitors with brain formation and neuro-development uncovered in IPA analysis ***(Supplementary figure 4H)***.

Based on the gene expression pattern of *Lyl-1*-deficient microglia and the signature of MΦ^Prim^ progenitors in the early YS, a contribution of *Lyl-1*-deficiency to neurodevelopmental disorders may be considered. Synaptic pruning and neural maturation, which characterize the perinatal phase of microglia development^*43*^, might be impaired in *Lyl-1*^*LacZ/LacZ*^ embryos considering the defects observed at P0-P3, the later key developmental stage regulated by Lyl-1. Indeed, *Lyl-1* deregulation has been observed in datasets reporting pathological models of brain myeloid cells (http://research-pub.gene.com/BrainMyeloidLandscape/#Mouse-gene/Mouse-gene/17095/geneReport.html)^*57*^, as well as in human neuropathies^*58, 59*^, including the 19p13.13 micro-deletion neuro-developmental disabilities.^*60*^ However, since Lyl-1 is expressed in endothelial cells, including in the brain^*61*^, a contribution of *LYL-1*-deficient endothelial cells to these diseases must be considered.

Altogether, our findings reveal Lyl-1 as a key factor regulating the production and differentiation of YS MΦ-progenitors and the development of microglia. Lyl-1 is the least studied partners of the transcription factor complex that regulates developmental hematopoiesis. The development of more appropriate models is required to precise Lyl-1 functions in microglia and determine its role in the development of other resident-MΦs populations.

## MATERIALS AND METHODS

### Mice

The following mouse strains were used: C57BL/6 (Charles Rivers Laboratories), called wild type (WT); *Lyl-1*^*LacZ,5*^; *Cx3cr1*^*GFP,62*^; *Cx3cr1*^*GFP/GFP*^*:Lyl-1*^*LacZ/LacZ*^ double knock-in strain (breeding schemes: ***supplemental information***). Experiments were conducted in compliance with French/European laws, under authorized project *#2016-030-5798*, approved by officially accredited local institutional animal (committee n°26) and French “Ministère de la Recherche” ethics boards.

### Tissues

E7.5-E10.5 YS and E10-FL were dissected as described.^*9, 63*^ For cytometry analyses, whole E9-E11 brains were prepared as described.^*22*^ After E12, microglia were recovered following Percoll (Sigma) separation, as described.^*64*^ The number of microglia per brain was estimated by reporting the percentage of CD11b^+^CD45^low^F4/80^+^ microglia to the cellularity of the corresponding sample.

### Tissue culture

YS explants, maintained for 1 day in organ culture in plates containing OptiMEM+Glutamax, 1% Penicillin-streptomycin, 0.1% β-mercaptoethanol (ThermoFisher) and 10% fetal calf serum (Hyclone), are referred to as OrgD1-YS. In clonogenic assays, YS suspension or sorted cells were plated in triplicate (3×10^3^ or 100-150 cells/mL) in Methocult® M3234 (StemCell Technologies) always supplemented the cytokines listed in ***supplemental table 5***. Colonies were scored at day 5 for primitive erythroblasts and day 7 for other progenitors.

### Flow cytometry

Cells, stained with antibodies listed in ***supplemental table 6***, for 30 min. on ice, were acquired (Canto II) or sorted (FACS-Aria III or Influx, BD Biosciences) and analyzed using FlowJo (Treestar) software.

The β-Galactosidase substrate fluorescein di-β-galactopyranoside (FDG; Molecular probe), was used as reporter for Lyl-1 expression in FACS-Gal assay.^*65,66*^ For apoptosis analysis, microglia were immune-stained and incubated with Annexin V-FITC. 7AAD was added before acquisition. For proliferation assays, pregnant females (12 gestational days) were injected with BrdU (10μM) and sacrificed 2 hours later. Microglia were isolated, immune-stained and prepared according to kit instruction (BD Pharmingen 552598). BrdU incorporation was revealed using anti-BrdU-APC.

### Brain imaging

To assess microglia morphology, the midbrain was dissected from *Cx3cr1*^*WT/GFP*^*:Lyl-1*^*WT/WT*^ and *Cx3cr1*^*WT/GFP*^*:Lyl-1*^*LacZ/LacZ*^ embryos, immune-labeled and image stacks collected using Leica SP8 confocal microscope ***(See Supplemental information)***.

### RT-qPCR analyses

RNA was extracted using Trizol. After cDNA synthesis (SuperScript™ VILO™ Master-Mix reverse transcriptase, ThermoFisher), Real Time (RT)-PCR was performed (SYBR Premix Ex TaqII, Takara Bio). Reference genes were *Actin, Hprt* and *Tubulin*. Gene expressions were normalized to the values obtained for E10-YS MΦ-progenitors, E12 WT Lin^-^Sca^+^cKit^+^ (LSK) progenitors or E12 WT microglia. Primers are listed in ***supplemental table 7***.

### RNA-seq

MΦ-progenitors (CD45^+^CD11b^+^Kit^+^) were sorted from E9 (MΦ^Prim^ progenitors) and E10 (MΦ^Prim^ + MΦ^T-Def^ progenitors) YS from WT or *Lyl-1*^*LacZ/LacZ*^ embryos. See ***supplemental information*** for sample processing, RNA-sequencing and analysis protocols. RNA-seq. data (accession number E-MTAB-9618) were deposited in EMBL-EBI ArrayExpress database (www.ebi.ac.uk/arrayexpress). Data were analyzed using Ingenuity® Pathway Analysis (IPA, QIAGEN)^*67*^, Gene set enrichment analysis (GSEA)^*68, 69*^, Morpheus and Venny softwares.

### Statistical analysis

Statistical tests were performed using Prism 7 (GraphPad). Statistical significance is indicated by p-values and/or as *p<0.05, **p<0.01, ***p<0.001 and ****p<0.0001.

## Acknowledgements

The authors thank Julien Bertrand for critical reading of the manuscript. We are grateful to the staff of the facilities at Gustave Roussy, the animal facility (PFEP, UMS AMMICa UMS 3655/US23, directed by P. Gonin), the imaging facility (PFIC, UMS AMMICa UMS 3655/US23, directed by C. Laplace-Builhe), the genomic facility directed by N. Droin, the bioinformatics facility (G. Meurice), directed by M. Deloger.

This work was supported by fundings from Institut National de la Santé et de la Recherche Médicale to W. Vainchenker, I. Plo and H. Raslova, from Centre National de la Recherche Scientifique and Université de Paris-Saclay to I. Godin, from grants INCA PLBio to I. Plo, “Ligue Nationale contre le Cancer” Certified Team to H. Raslova, “Association pour la Recherche sur le Cancer” (n°4878) to I. Godin, Gustave Roussy (TA DERE 17) to D. Ren, Grant Agency of the Czech Republic (GACR n°19–23154S) to D. Filipp and from fellowships from “Association pour la Recherche sur le Cancer” to A.-L. Kaushik; “Société Française d’Hématologie” to S. Wang and Chinese Scholarship Council fellowships to S. Wang and D. Ren.

## Author Information

The authors declare no competing financial interests.

Correspondence should be addressed to I.G.

## REFERENCES

1. Pina, C. & Enver, T. Differential contributions of haematopoietic stem cells to foetal and adult haematopoiesis: insights from functional analysis of transcriptional regulators. Oncogene 26, 6750–6765 (2007).

2. Wilson, N.K. et al. Combinatorial Transcriptional Control In Blood Stem/Progenitor Cells: Genome-wide Analysis of Ten Major Transcriptional Regulators. Cell Stem Cell 7, 532–544 (2010).

3. Curtis, D.J., Salmon, J.M. & Pimanda, J.E. Concise Review: Blood Relatives: Formation and regulation of hematopoietic stem cells by the basic helix-loop-helix transcription factors stem cell leukemia and lymphoblastic leukemia-derived sequence 1. Stem Cells 30, 1053–1058 (2012).

4. Porcher, C., Chagraoui, H. & Kristiansen, M.S. SCL/TAL1: a multifaceted regulator from blood development to disease. Blood 129, 2051–2060 (2017).

5. Capron, C. et al. The SCL relative LYL-1 is required for fetal and adult hematopoietic stem cell function and B-cell differentiation. Blood 107, 4678–4686 (2006).

6. McGrath, K.E., Frame, J.M. & Palis, J. Early hematopoiesis and macrophage development. Seminars in Immunology 27, 379–387 (2015).

7. Palis, J. Hematopoietic stem cell-independent hematopoiesis: emergence of erythroid, megakaryocyte, and myeloid potential in the mammalian embryo. FEBS Lett 590, 3965–3974 (2016).

8. Cumano, A. & Godin, I. Ontogeny of the hematopoietic system. Annu Rev Immunol 25, 745–785 (2007).

9. Kieusseian, A., Brunet de la Grange, P., Burlen-Defranoux, O., Godin, I. & Cumano, A. Immature hematopoietic stem cells undergo maturation in the fetal liver. Development 139, 3521–3530 (2012).

10. Palis, J., Robertson, S., Kennedy, M., Wall, C. & Keller, G. Development of erythroid and myeloid progenitors in the yolk sac and embryo proper of the mouse. Development 126, 5073–5084 (1999).

11. Bertrand, J.Y. et al. Three pathways to mature macrophages in the early mouse yolk sac. Blood 106, 3004–3011 (2005).

12. Tober, J. et al. The megakaryocyte lineage originates from hemangioblast precursors and is an integral component both of primitive and of definitive hematopoiesis. Blood 109, 1433–1441 (2007).

13. Sumner, R., Crawford, A., Mucenski, M. & Frampton, J. Initiation of adult myelopoiesis can occur in the absence of c-Myb whereas subsequent development is strictly dependent on the transcription factor. Oncogene 19, 3335–3342 (2000).

14. Schulz, C. et al. A lineage of myeloid cells independent of Myb and hematopoietic stem cells. Science 336, 86–90 (2012).

15. Hoeffel, G. & Ginhoux, F. Fetal monocytes and the origins of tissue-resident macrophages. Cell Immunol 330, 5–15 (2018).

16. Utz, S.G. et al. Early Fate Defines Microglia and Non-parenchymal Brain Macrophage Development. Cell 181, 557–573 e518 (2020).

17. Ginhoux, F. & Guilliams, M. Tissue-Resident Macrophage Ontogeny and Homeostasis. Immunity 44, 439–449 (2016).

18. Mass, E. Delineating the origins, developmental programs and homeostatic functions of tissue-resident macrophages. Int Immunol 30, 493–501 (2018).

19. Ginhoux, F. et al. Fate Mapping Analysis Reveals That Adult Microglia Derive from Primitive Macrophages. Science 330, 841–845 (2010).

20. Gomez Perdiguero, E. et al. Tissue-resident macrophages originate from yolk-sac-derived erythro-myeloid progenitors. Nature 518, 547–551 (2015).

21. Kierdorf, K. et al. Microglia emerge from erythromyeloid precursors via Pu.1-and Irf8-dependent pathways. Nat Neurosci 16, 273–280 (2013).

22. Alliot, F., Godin, I. & Pessac, B. Microglia derive from progenitors, originating from the yolk sac, and which proliferate in the brain. Brain Res Dev Brain Res 117, 145–152 (1999).

23. Wu, Y. & Hirschi, K.K. Tissue-Resident Macrophage Development and Function. Frontiers in Cell and Developmental Biology 8 (2021).

24. Herbomel, P., Thisse, B. & Thisse, C. Zebrafish early macrophages colonize cephalic mesenchyme and developing brain, retina, and epidermis through a M-CSF receptor-dependent invasive process. Dev Biol 238, 274–288 (2001).

25. Ferrero, G. et al. Embryonic Microglia Derive from Primitive Macrophages and Are Replaced by cmyb-Dependent Definitive Microglia in Zebrafish. Cell Reports 24, 130–141 (2018).

26. Wittamer, V. & Bertrand, J.Y. Yolk sac hematopoiesis: does it contribute to the adult hematopoietic system? Cell Mol Life Sci (2020).

27. Azzoni, E. et al. Kit ligand has a critical role in mouse yolk sac and aorta-gonad-mesonephros hematopoiesis. EMBO reports (2018).

28. Espin-Palazon, R. et al. Proinflammatory signaling regulates hematopoietic stem cell emergence. Cell 159, 1070–1085 (2014).

29. Giroux, S. et al. lyl-1 and tal-1/scl, two genes encoding closely related bHLH transcription factors, display highly overlapping expression patterns during cardiovascular and hematopoietic ontogeny. Gene Expr Patterns 7, 215–226 (2007).

30. Cumano, A., Dieterlen-Lievre, F. & Godin, I. Lymphoid potential, probed before circulation in mouse, is restricted to caudal intraembryonic splanchnopleura. Cell 86, 907–916 (1996).

31. Cumano, A., Ferraz, J.C., Klaine, M., Di Santo, J.P. & Godin, I. Intraembryonic, but not yolk sac hematopoietic precursors, isolated before circulation, provide long-term multilineage reconstitution. Immunity 15, 477–485 (2001).

32. Hoeffel, G. et al. C-Myb+ Erythro-Myeloid Progenitor-Derived Fetal Monocytes Give Rise to Adult Tissue-Resident Macrophages. Immunity 42, 665–678 (2015).

33. Mass, E. et al. Specification of tissue-resident macrophages during organogenesis. Science 353 (2016).

34. Espin-Palazon, R., Weijts, B., Mulero, V. & Traver, D. Proinflammatory Signals as Fuel for the Fire of Hematopoietic Stem Cell Emergence. Trends in Cell Biology 28, 58–66 (2018).

35. Mariani, S.A. et al. Pro-inflammatory Aorta-Associated Macrophages Are Involved in Embryonic Development of Hematopoietic Stem Cells. Immunity (2019).

36. Palpant, N.J. et al. Chromatin and Transcriptional Analysis of Mesoderm Progenitor Cells Identifies HOPX as a Regulator of Primitive Hematopoiesis. Cell Reports 20, 1597–1608 (2017).

37. Chiu, S.K. et al. Shared roles for Scl and Lyl1 in murine platelet production and function. Blood 134, 826–835 (2019).

38. McCormack, M.P. et al. Requirement for Lyl1 in a model of Lmo2-driven early T-cell precursor ALL. Blood 122, 2093–2103 (2013).

39. Epelman, S. et al. Embryonic and Adult-Derived Resident Cardiac Macrophages Are Maintained through Distinct Mechanisms at Steady State and during Inflammation. Immunity 40, 91–104 (2014).

40. Leid, J. et al. Primitive Embryonic Macrophages are Required for Coronary Development and Maturation. Circulation Research 118, 1498 (2016).

41. Hagemeyer, N. et al. Transcriptome-based profiling of yolk sac-derived macrophages reveals a role for Irf8 in macrophage maturation. EMBO J 35, 1730–1744 (2016).

42. Ferrier, R. et al. Physical interaction of the bHLH LYL1 protein and NF-kappaB1 p105. Oncogene 18, 995–1005 (1999).

43. Matcovitch-Natan, O. et al. Microglia development follows a stepwise program to regulate brain homeostasis. Science 353, aad8670 (2016).

44. Goldmann, T. et al. Origin, fate and dynamics of macrophages at central nervous system interfaces. Nature Immunology 17, 797–805 (2016).

45. Lavin, Y. et al. Tissue-Resident Macrophage Enhancer Landscapes Are Shaped by the Local Microenvironment. Cell 159, 1312–1326 (2014).

46. Zhang, Y. et al. An RNA-sequencing transcriptome and splicing database of glia, neurons, and vascular cells of the cerebral cortex. J Neurosci 34, 11929–11947 (2014).

47. Zeisel, A. et al. Brain structure. Cell types in the mouse cortex and hippocampus revealed by single-cell RNA-seq. Science 347, 1138–1142 (2015).

48. Wehrspaun, C.C., Haerty, W. & Ponting, C.P. Microglia recapitulate a hematopoietic master regulator network in the aging human frontal cortex. Neurobiol Aging 36, 2443 e2449–2443 e2420 (2015).

49. Bennett, M.L. et al. New tools for studying microglia in the mouse and human CNS. Proceedings of the National Academy of Sciences 113, E1738–E1746 (2016).

50. Zhan, Y. et al. Deficient neuron-microglia signaling results in impaired functional brain connectivity and social behavior. Nat Neurosci 17, 400–406 (2014).

51. Paolicelli, R.C. et al. Synaptic pruning by microglia is necessary for normal brain development. Science 333, 1456–1458 (2011).

52. Antony, J.M., Paquin, A., Nutt, S.L., Kaplan, D.R. & Miller, F.D. Endogenous microglia regulate development of embryonic cortical precursor cells. J Neurosci Res 89, 286–298 (2011).

53. Ueno, M. et al. Layer V cortical neurons require microglial support for survival during postnatal development. Nat Neurosci 16, 543–551 (2013).

54. Soucie, E.L. et al. Lineage-specific enhancers activate self-renewal genes in macrophages and embryonic stem cells. Science 351 (2016).

55. Crotti, A. & Ransohoff, R.M. Microglial Physiology and Pathophysiology: Insights from Genome-wide Transcriptional Profiling. Immunity 44, 505–515 (2016).

56. Bisht, K., Sharma, K. & Tremblay, M.-È. Chronic stress as a risk factor for Alzheimer’s disease: Roles of microglia-mediated synaptic remodeling, inflammation, and oxidative stress. Neurobiology of Stress 9, 9–21 (2018).

57. Friedman, B.A. et al. Diverse Brain Myeloid Expression Profiles Reveal Distinct Microglial Activation States and Aspects of Alzheimer’s Disease Not Evident in Mouse Models. Cell Reports 22, 832–847 (2018).

58. Colangelo, V. et al. Gene expression profiling of 12633 genes in Alzheimer hippocampal CA1: transcription and neurotrophic factor down-regulation and up-regulation of apoptotic and pro-inflammatory signaling. J Neurosci Res 70, 462–473 (2002).

59. Thomas, D.M., Francescutti-Verbeem, D.M. & Kuhn, D.M. Gene expression profile of activated microglia under conditions associated with dopamine neuronal damage. Faseb J 20, 515–517 (2006).

60. Nimmakayalu, M. et al. Apparent germline mosaicism for a novel 19p13.13 deletion disrupting NFIX and CACNA1A. Am J Med Genet A 161A, 1105-1109 (2013).

61. Pirot, N. et al. LYL1 activity is required for the maturation of newly formed blood vessels in adulthood. Blood 115, 5270–5279 (2010).

62. Jung, S. et al. Analysis of fractalkine receptor CX(3)CR1 function by targeted deletion and green fluorescent protein reporter gene insertion. Mol Cell Biol 20, 4106–4114 (2000).

63. Bertrand, J.Y., Giroux, S., Cumano, A. & Godin, I. Hematopoietic stem cell development during mouse embryogenesis. Methods Mol Med 105, 273–288 (2005).

64. Mildner, A. et al. Microglia in the adult brain arise from Ly-6ChiCCR2+ monocytes only under defined host conditions. Nat Neurosci 10, 1544–1553 (2007).

65. Fiering, S.N. et al. Improved FACS-Gal: flow cytometric analysis and sorting of viable eukaryotic cells expressing reporter gene constructs. Cytometry 12, 291–301 (1991).

66. Guo, W. & Wu, H. Detection of LacZ expression by FACS-Gal analysis. Nature Protocol Exchange (2008).

67. Krämer, A., Green, J., Pollard, J., Jr. & Tugendreich, S. Causal analysis approaches in Ingenuity Pathway Analysis. Bioinformatics 30, 523–530 (2014).

68. Mootha, V.K. et al. PGC-1alpha-responsive genes involved in oxidative phosphorylation are coordinately downregulated in human diabetes. Nat Genet 34, 267–273 (2003).

69. Subramanian, A. et al. Gene set enrichment analysis: a knowledge-based approach for interpreting genome-wide expression profiles. Proc Natl Acad Sci U S A 102, 15545–15550 (2005).

