## Supplemental data for "Lyl-1 regulates primitive macrophages and microglia development"

#### *SUPPLEMENTAL INFORMATION*

##### *Supplemental material and methods*

**Mice.**

The following mouse strains, housed in Gustave Roussy Institute animal facility (License #H94-076-11) were used: 1- C57BL/6 mice (Harlan or Charles Rivers Laboratories), referred to as wild type (WT); 2- *Lyl-1*$^{LacZ}$ mice.[1] To avoid the possible detection of FDG/Lyl-1 expression from maternally-derived M$\Phi$[2] in heterozygous embryos, *Lyl-1*$^{LacZ/LacZ}$ males were crossed with WT females; 3- *Cx3cr1*$^{GFP}$ mice.[3] *Cx3cr1*$^{GFP/GFP}$ males were crossed with WT females to generate *Cx3cr1*$^{WT/GFP}$ mice/embryos or *Lyl-1*$^{LacZ/LacZ}$ females to generate *Cx3cr1*$^{WT/GFP}$*:Lyl-1*$^{WT/LacZ}$ mice/embryos. 4- *Cx3cr1*$^{GFP/GFP}$*:Lyl-1*$^{LacZ/LacZ}$ double mutant strain developed from *Cx3cr1*$^{WT/GFP}$*:Lyl-1*$^{WT/LacZ}$ crosses. *Cx3cr1*$^{WT/GFP}$*:Lyl-1*$^{WT/LacZ}$ and *Cx3cr1*$^{WT/GFP}$*:Lyl-1*$^{LacZ/LacZ}$ mice/embryos were obtained by crossing *Cx3cr1*$^{GFP/GFP}$*:Lyl-1*$^{LacZ/LacZ}$ males respectively to C57BL/6 or *Lyl-1*$^{LacZ/LacZ}$ females. Experiments were conducted in compliance with French/European laws (project *#2016-030-5798)*.

The day of vaginal plug observation was considered as E0.5. Pregnant females were sacrificed by cervical dislocation. Pre-somite embryos were staged according to Downs et al.[4], by somite counting from E8 to E10.5 and thereafter by morphological landmarks.

**Brain imaging information, related to figure 4**

To assess microglia morphology in E12 embryos, the midbrain was dissected from *Cx3cr1*$^{WT/GFP}$*:Lyl-1*$^{WT/WT}$ and *Cx3cr1*$^{WT/GFP}$*:Lyl-1*$^{LacZ/LacZ}$ embryos and sectioned through the midline. After fixation (4% paraformaldehyde) overnight at 4°C, whole midbrains were washed in phosphate-buffered saline (PBS)/0.1M glycine and incubated overnight in PBS/15% sucrose at 4°C. Midbrains were washed with PBS+0.1% Tween and incubated 90min. in blocking buffer (PBS+10% FCS) at room temperature (RT). Midbrains were subsequently immune-labeled with F4/80-APC overnight at 4°C. After washing, they were incubated 3 min. in PBS+DAPI (1 µg/mL) at RT and washed. Finally, midbrains were placed in the central well of glass-bottom culture dishes (P35G-1.5-10-C; MatTek, USA) filled with PBS+10% FCS. After appropriate orientation of the sample, the well was covered with a 12mm ∅ glass coverslip. Image stacks were collected using a Leica SP8 confocal microscope. To unsure an unbiased choice of the cells imaged, taking into account possible changes in cell distribution induced by *Lyl-1* deficiency, we always acquired cells in similar positions regarding the landmark set in the midbrain flat mount, as shown in ***Supplemental data and figure 4C***. Images were processed using Imaris x64 (version 7.7.2; Bitplane) and Photoshop 8.0 (Adobe Systems, San Jose, CA) softwares.

**RNA-seq.**

***Sample preparation:*** MΦP (Kit$^+$CD45$^+$CD11b$^+$) were sorted from E9 (MΦ$^{Prim}$ progenitor) and E10 (MΦ$^{Prim}$ + MΦ$^{T\text{-}Def}$ progenitors) YS pools from WT or *Lyl-1*$^{LacZ/LacZ}$ embryos (Four biological replicates). RNA was extracted as described above.

***Sample processing:*** The RNA integrity (RNA Integrity Score≥7.0) was checked on the Agilent 2100 Bioanalyzer (Agilent) and quantity was determined using Qubit (Invitrogen). SureSelect Automated Strand Specific RNA Library Preparation Kit was used according to manufacturer's instructions with the Bravo Platform. Briefly, 50ng of total RNA sample was used for poly-A mRNA selection using oligo(dT) beads and subjected to thermal mRNA fragmentation. Fragmented mRNA samples were subjected to cDNA synthesis and converted into double stranded DNA using reagents supplied in the kit. The resulting dsDNA was used for library preparation. The final libraries were bar-coded, purified, pooled together in equal concentrations and subjected to paired-end sequencing (2x100) on Novaseq-6000 sequencer (Illumina) at Gustave Roussy genomic facility. RNA-seq. data (accession number E-MTAB-9618) were deposited in the ArrayExpress database at EMBL-EBI (www.ebi.ac.uk/arrayexpress).

***RNA-seq. analysis:*** Quality of RNA-seq. reads was assessed with FastQC 0.11.7 and MultiQC 1.5.[5] Low quality reads were trimmed with Trimmomatic 0.33.[6] Salmon 0.9.0 tool[7] was used for quantifying the expression of transcripts using geneset annotation from Gencode project release M17 for mouse.[8] The version of transcriptome reference sequences used was GRCm38.p6.

Statistical analysis was performed using R with the method proposed by Anders and Huber implemented in the DESeq2 Bioconductor package.[9] The differential expression analysis in DESeq2 uses a generalized linear model (GLM) where counts are modeled using a negative binomial distribution. Counts were normalized from the estimated size factors using the median ratio method and a Wald test was used to test the significance of GLM coefficients. Genes were considered differentially expressed when adjusted p-value < 0.05 and fold-change > 2.

Data were analyzed through the use of Ingenuity® Pathway Analysis (QIAGEN Inc., https://www.qiagenbioinformatics.com/products/ingenuity-pathway-analysis)[10], Gene set enrichment analysis (GSEA; https://www.gsea-msigdb.org/gsea/index.jsp)[11,12], Morpheus (https://software.broadinstitute.org/morpheus/) and Venny (https://bioinfogp.cnb.csic.es/tools/venny/) softwares.

***Supplemental information related to Figure 1C-E, 3A and Supplemental figure 2A-D***

During YS development, both $M\Phi^{Prim}$ and $M\Phi^{T\text{-}Def}$ progenitors originate from $cKit^{+}CD31^{+}CD45^{-}$ progenitors (C subset), which differentiate into MΦs via 3 subsets (A1 to A3)[2] ***(Supplemental figure 2A)***. In *Lyl-1*$^{LacZ/LacZ}$ E9-YS, a large $CD11b^{+}CD31^{+}CD45^{neg/low}$ subset was identified through flow cytometry analyses of the distribution of MΦ progenitor subsets. This $CD11b^{+}CD31^{+}CD45^{neg/low}$ subset was nearly absent from WT E9-YS and it disappeared from mutant YS after E9.5

***(Supplemental figure 2C)***. This transient subset, which display a phenotype intermediate between the C and A1 subsets, was also characterized by the expression of Lyl-1, as shown using Facs-Gal assay ***(Supplemental figure 2D)***. Together with the increased production of $M\Phi^{Prim}$ progenitors observed in clonogenic assays in mutant E8-YS (***Figure 3A)***, this observation suggests that in a WT context, Lyl-1 might negatively regulate the commitment, expansion and/or timing of production of $M\Phi^{Prim}$.

***Supplemental figures***

***Supplemental figure 1, related to figure 1:***

**A.** Gating strategy used to analyze $FDG^{+}$/Lyl-$1^{+}$ expression in MΦ progenitors from E9- and E9.5-YS from WT, *Lyl-1*$^{WT/LacZ}$ and *Lyl-1*$^{LacZ/LacZ}$ embryos ***(Figure 1B)*** and in MΦ-progenitors and mature MΦ-progenitors from E10 WT and *Lyl-1*$^{LacZ/LacZ}$ YS, FL and brain ***(Figure 4A, 5A, C, G)***. Shown is WT YS sample at E10.

**B.** Gene set enrichment analysis (GSEA) of the whole transcriptome of WT E9-YS MΦ-progenitors, compared to EMP (left) and MΦs (right) signatures defined by Mass *et al.*[13] (NES: normalized enrichment score; FDR: false discovery rate).

**C.** Top 1 IPA network representing the molecular relationships between DEG in E9 vs. E10 WT MΦ-progenitors indicated an overexpression of MHC-II at E9, while E10 progenitors over-expressed the TNFα signaling pathway.

**D.** Flow cytometry analysis of MHC-II expression in MΦ-progenitors from WT E9- and E10-YS. MHC-II expression is significantly reduced in E10 $Kit^{+}CD45^{+}CD11b^{+}$ MΦ-progenitors compared to E9, irrelevant of the genotype.

**E.** Top 5 IPA network Illustrates the bias towards inflammatory signaling in E10 WT MΦ-progenitors, compared to those present at E9.

**F.** WT MΦ-progenitors at E10 were enriched in TLR and TNFα signaling pathways (GSEA pathways).

**G.** WT MΦ-progenitors at E10 were enriched in TGFβ signaling pathway (GSEA pathways).

**H.** WT MΦ-progenitors at E10 were enriched in genes belonging to the complement cascade (KEGG pathway).

**I.** Expression profiles of DEGs related to phagocytosis (Heatmap displays transformed log2-expression values; unpaired *t*-Test, two-tailed).

***Supplemental figure 2, related to figure 3:***

**A. Phenotype of MΦ subsets.** MΦ develop from $Kit^{+}CD31^{+}CD45^{-}$ progenitors (C subset). MΦ-progenitors first acquire the expression of CD11b and CD45 (A1 subset), then down-regulate Kit expression (A2 subset). The differentiation to mature MΦs is hallmarked by the concomitant down-regulation of CD31 and up-regulation of F4/80 and $CX_3CR1$ (A3 subset).

**B.** Gating strategy used to characterize MΦ-progenitors development from E9- and E9.5-YS. Note that this gating strategy differs from the one used to analyze the E9-10 MΦs ***(Supplemental figure 1A)*** in the YS and brain, which lack the intermediate progenitor subset ($CD11b^{+}CD31^{+}CD45^{neg/low}$).

**C. Lyl-1-deficiency leads to an increased commitment of mesodermal/pre-hematopoietic cells to a MΦ fate.** At E9, *Lyl-1*$^{WT/LacZ}$ and *Lyl-1*$^{LacZ/LacZ}$ YS harbor a large $CD45^{neg/low}$ subset within CD11b+ CD31+ cells (% in top left profile), which nearly absent from WT YS samples. This C to A1 transition subset is no longer present in mutant YS at E9.5 (lower profiles). Right panel: Quantification of the C to A1 transition subset.

**D. Facs-Gal analysis of Lyl-1 expression in E9-YS MΦ-progenitors.** The contour plots in WT samples indicate the level of non-specific β-Gal activity. Both $CD45^{+}$ and the C to A1/CD45- subsets within the $CD11b^{+}$ $CD31^{+}$ MΦP gate (% in top left profile) express FDG/Lyl-1.

**E.** Enrichment plot of top 1 GSEA pathway down-regulated in *Lyl-1*$^{LacZ/LacZ}$ MΦ-progenitors compared to WT at E10 is related to heart development (NES: Normalized enrichment score; FDR: False discovery rate).

**F. Lyl-1 deficiency leads to an increased perinatal lethality:** The number of *Lyl-1*$^{LacZ/LacZ}$ embryos per litter was unmodified at E14 (left), but the number of living newborns (middle) was reduced. The number of *Lyl-1*$^{LacZ/LacZ}$ pups was further reduced at weaning (right) with a significant increase of neonatal lethality until

P15 (main occurrence between P1 and P5: data not shown). *Lyl-1*$^{LacZ/LacZ}$ had a balanced sex ratio at birth and weaning (data not shown). Similar numbers of E14 embryos, newborns and weanling were obtained from WT and *Lyl-1*$^{WT/LacZ}$ resulting from WT x *Lyl-1*$^{LacZ/LacZ}$ crosses (E14: n = WT: 41; *Lyl-1*$^{WT/LacZ}$: 39; *Lyl-1*$^{LacZ/LacZ}$: 33. Newborns and weanlings: n = WT: 230; *Lyl-1*$^{WT/LacZ}$: 95; *Lyl-1*$^{LacZ/LacZ}$: 432 (Tuckey box plot; Unpaired, two-tailed *t*-Test).

***Supplemental figure 3, related to Figure 4***

**A. RT-qPCR analyses of *Lyl-1* expression in A1 to A3 MΦ subsets from *Cx3cr1*$^{WT/GFP}$ E10-YS**. *Lyl-1* is expressed by the 3 subsets with an expression level decreasing upon differentiation. Expression levels were normalized to the mean value obtained for *Cx3cr1*$^{WT/GFP}$ A1 progenitors (n=3).

**B.** Top GSEA pathways weakened in *Lyl-1*$^{LacZ/LacZ}$ compared to WT MΦ-progenitors at E9 (FDR q-value <0.05). Highlighted are pathways related to cytokine signaling (green), pathways enriched in E9 WT compared to E10 WT that are down-regulated due to *Lyl-1*-deficiency (orange) and the pathways enriched in E10 WT compared to E9 WT that were further reduced in *Lyl-1*$^{LacZ/LacZ}$ MΦ-progenitors at E9 (purple).

**C.** Left: *Spi1*/PU.1 (Top 2 GSEA TF pathway) signaling was weaker in E9 *Lyl-1*$^{LacZ/LacZ}$ than in WT E9 MΦ-progenitors (NES: normalized enrichment score; FDR: false discovery rate). Right: Expression profiles of DEGs related to *Spi1*/PU.1 pathway (Heatmap displays transformed log2-expression values; unpaired t-Test, two-tailed).

**D.** Left: The top 1 GO enrichment plot pointed to a defective inflammatory response in E9 *Lyl-1*$^{LacZ/LacZ}$ MΦ-progenitors (NES: normalized enrichment score; FDR: false discovery rate). Right: Expression profiles of DEGs related to inflammatory response and cytokine signaling. (Heatmap displays transformed log2-expression values; Unpaired *t*-Test, two-tailed).

**E.** NFkB (left: Top1 KEGG pathway) and Toll-like receptor (right: Top 2 GSEA pathway) signaling were weaker in E9 *Lyl-1*$^{LacZ/LacZ}$, compared to WT E9 MΦ-progenitors (NES: normalized enrichment score; FDR: false discovery rate).

***Supplemental figure 4, related to figure 6***

**A. Microglia express Lyl-1 from embryonic stages to adulthood.** FDG/Lyl expression in WT and *Lyl-1*$^{LacZ/LacZ}$ brain from E12 to adult stages. In *Lyl-1*$^{LacZ/LacZ}$ mutants, the entire microglia population is FDG$^+$/Lyl-1$^+$, but Lyl-1 expression level decreases in the adult, with a clear transition occurring at P15. Plain grey histograms indicate non-specific background $\beta$-Gal activity/FDG levels in WT samples.

**B.** Annexin V-7AAD quantification of apoptosis levels in CD11b$^+$F4/80$^+$CD45$^{low}$ microglia at E12 and E14. No significant genotype-associated modification of microglia apoptosis level was observed in E12 microglia (right). At E14 (left), the apoptosis level was significantly lower in *Lyl-1*$^{LacZ/LacZ}$ compared to WT and *Lyl-1*$^{WT/LacZ}$ brains, which may account for the recovery of microglia pool size after E12.

**C.** The midbrain was dissected from *Cx3cr1*$^{WT/GFP}$*:Lyl-1*$^{WT/WT}$ and *Cx3cr1*$^{WT/GFP}$*:Lyl-1*$^{LacZ/LacZ}$ embryos (left). After removal of the meningeal layer (M), the midbrain was sectioned along the midline (middle left) and flat mounted (middle right). To unsure an unbiased choice, cells were always acquired in the same location (right) using the midline (arrowhead) and lateral thickening as landmarks: the cell closest to the middle of the frame (arrow) was acquired (Bar=1mm).

**D.** Confocal imaging of the midbrain of *Cx3cr1*$^{WT/GFP}$ and *Cx3cr1*$^{WT/GFP}$*:Lyl-1*$^{LacZ/LacZ}$ embryos at E12 pointed to *Lyl-1*$^{LacZ/LacZ}$-induced morphological changes in microglia. Microglia were identified by *Cx3cr1*-driven GFP expression and F4/80 immunostaining (Bar=100 $\mu$m).

**E.** *Lyl-1*-deficiency does not lead to modifications of cells recovery from E9 to E14 whole brains.

**F.** Cytometry analyses evidenced no genotype-associated difference in microglia pool size in the brain from adult WT and *Lyl-1*$^{WT/LacZ}$ mice.

**G.**: CD11b$^+$F4/80$^+$CD45$^{low}$ microglia was isolated at sequential development stages for quantitative RT-PCR analyses. Bar graphs show the kinetic of expression of a selected set of genes that were not significantly modified in *Lyl-1* mutants at E12 and P0-P3, but may be so at post-natal stages. Gene expressions were normalized to the mean expression value in WT E12 microglia (n=3). Error bars indicate s.e.m. (*P* values were determined by unpaired, two-tailed *t*-Test).

**H.** IPA function prediction: A function in neurogenesis is predicted for E9 *Lyl-1*$^{LacZ/LacZ}$ M$\Phi$-progenitors considering the DEGs enriched compared to E9 WT.

#### *References*


1. Capron C, Lécluse Y, Kaushik AL, et al. The SCL relative LYL-1 is required for fetal and adult hematopoietic stem cell function and B-cell differentiation. *Blood*. 2006;107(12):4678-4686.
2. Bertrand JY, Jalil A, Klaine M, Jung S, Cumano A, Godin I. Three pathways to mature macrophages in the early mouse yolk sac. *Blood*. 2005;106(9):3004-3011.
3. Jung S, Aliberti J, Graemmel P, et al. Analysis of fractalkine receptor CX(3)CR1 function by targeted deletion and green fluorescent protein reporter gene insertion. *Mol Cell Biol*. 2000;20(11):4106-4114.
4. Downs KM, Davies T. Staging of gastrulating mouse embryos by morphological landmarks in the dissecting microscope. *Development*. 1993;118:1255-1266.
5. Ewels P, Magnusson M, Lundin S, Käller M. MultiQC: summarize analysis results for multiple tools and samples in a single report. *Bioinformatics*. 2016;32(19):3047-3048.
6. Bolger AM, Lohse M, Usadel B. Trimmomatic: a flexible trimmer for Illumina sequence data. *Bioinformatics*. 2014;30(15):2114-2120.
7. Patro R, Duggal G, Love MI, Irizarry RA, Kingsford C. Salmon provides fast and bias-aware quantification of transcript expression. *Nat Methods*. 2017;14(4):417-419.
8. Frankish A, Diekhans M, Ferreira AM, et al. GENCODE reference annotation for the human and mouse genomes. *Nucleic Acids Res*. 2019;47(D1):D766-d773.
9. Love MI, Huber W, Anders S. Moderated estimation of fold change and dispersion for RNA-seq data with DESeq2. *Genome Biol*. 2014;15(12):550.
10. Krämer A, Green J, Pollard J, Jr., Tugendreich S. Causal analysis approaches in Ingenuity Pathway Analysis. *Bioinformatics*. 2014;30(4):523-530.
11. Mootha VK, Lindgren CM, Eriksson KF, et al. PGC-1alpha-responsive genes involved in oxidative phosphorylation are coordinately downregulated in human diabetes. *Nat Genet*. 2003;34(3):267-273.
12. Subramanian A, Tamayo P, Mootha VK, et al. Gene set enrichment analysis: a knowledge-based approach for interpreting genome-wide expression profiles. *Proc Natl Acad Sci U S A*. 2005;102(43):15545-15550.
13. Mass E, Ballesteros I, Farlik M, et al. Specification of tissue-resident macrophages during organogenesis. *Science*. 2016;353(6304).

### Supplemental figure 1

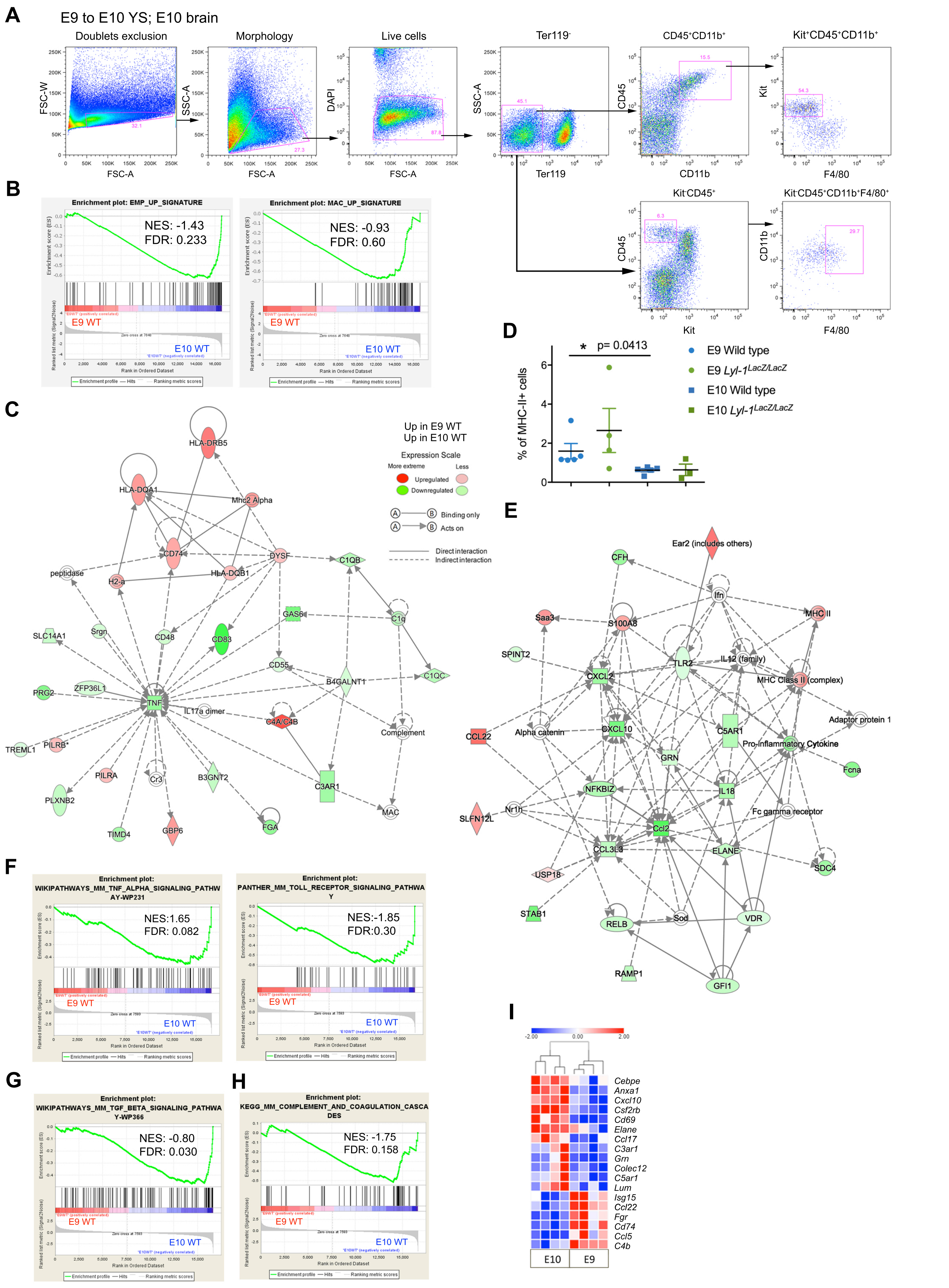

Supplemental Figure 2

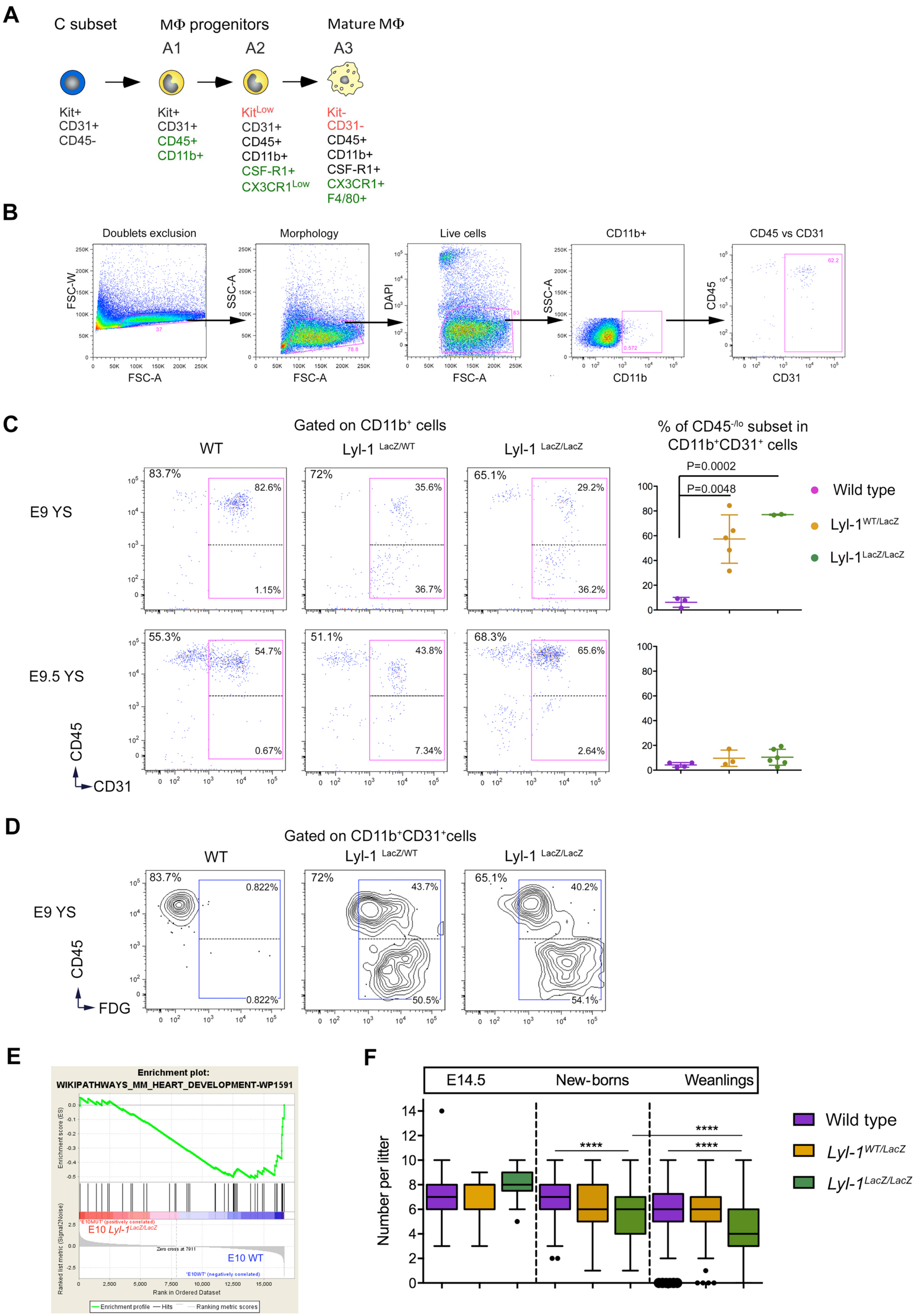

Supplemental figure 3

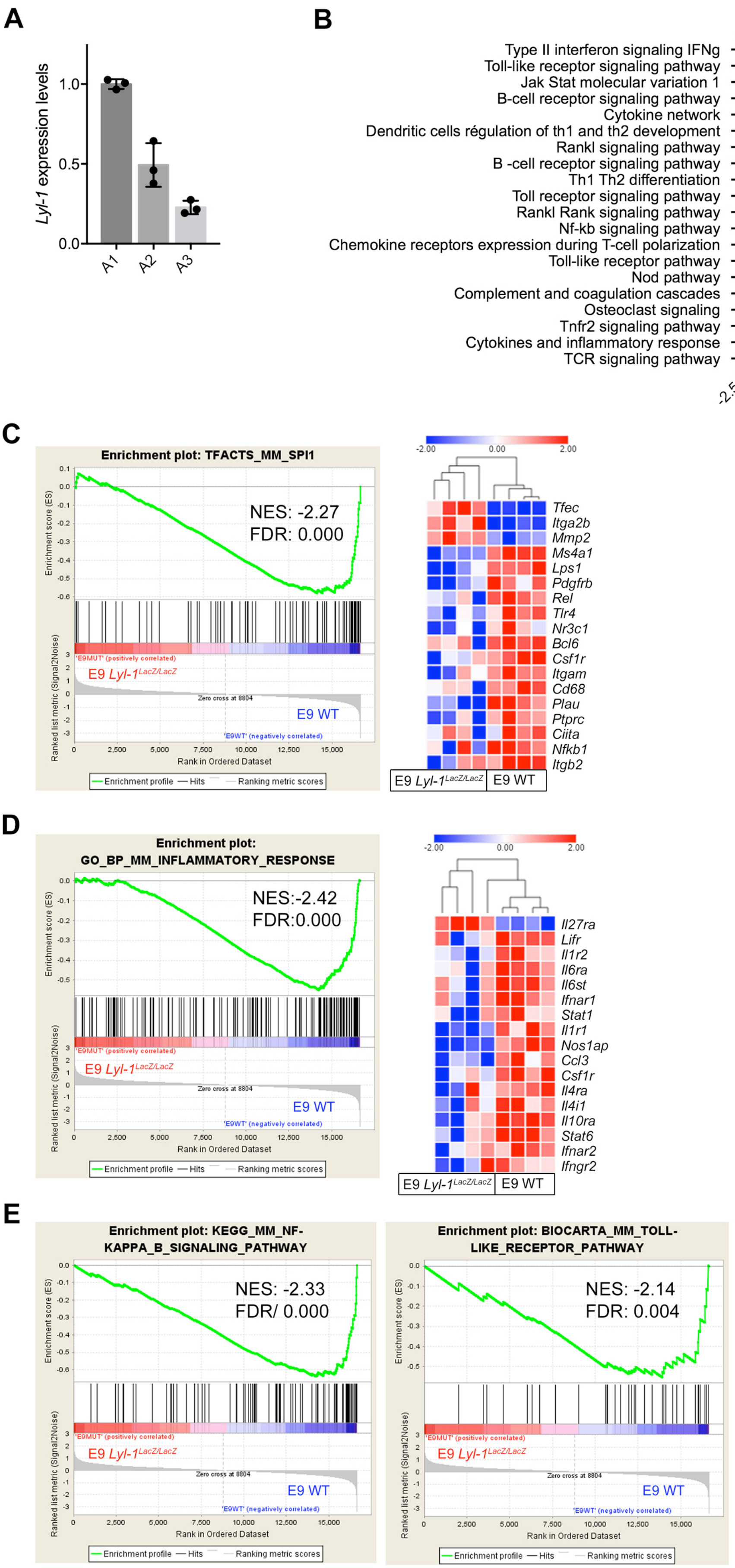

Supplemental Figure 4

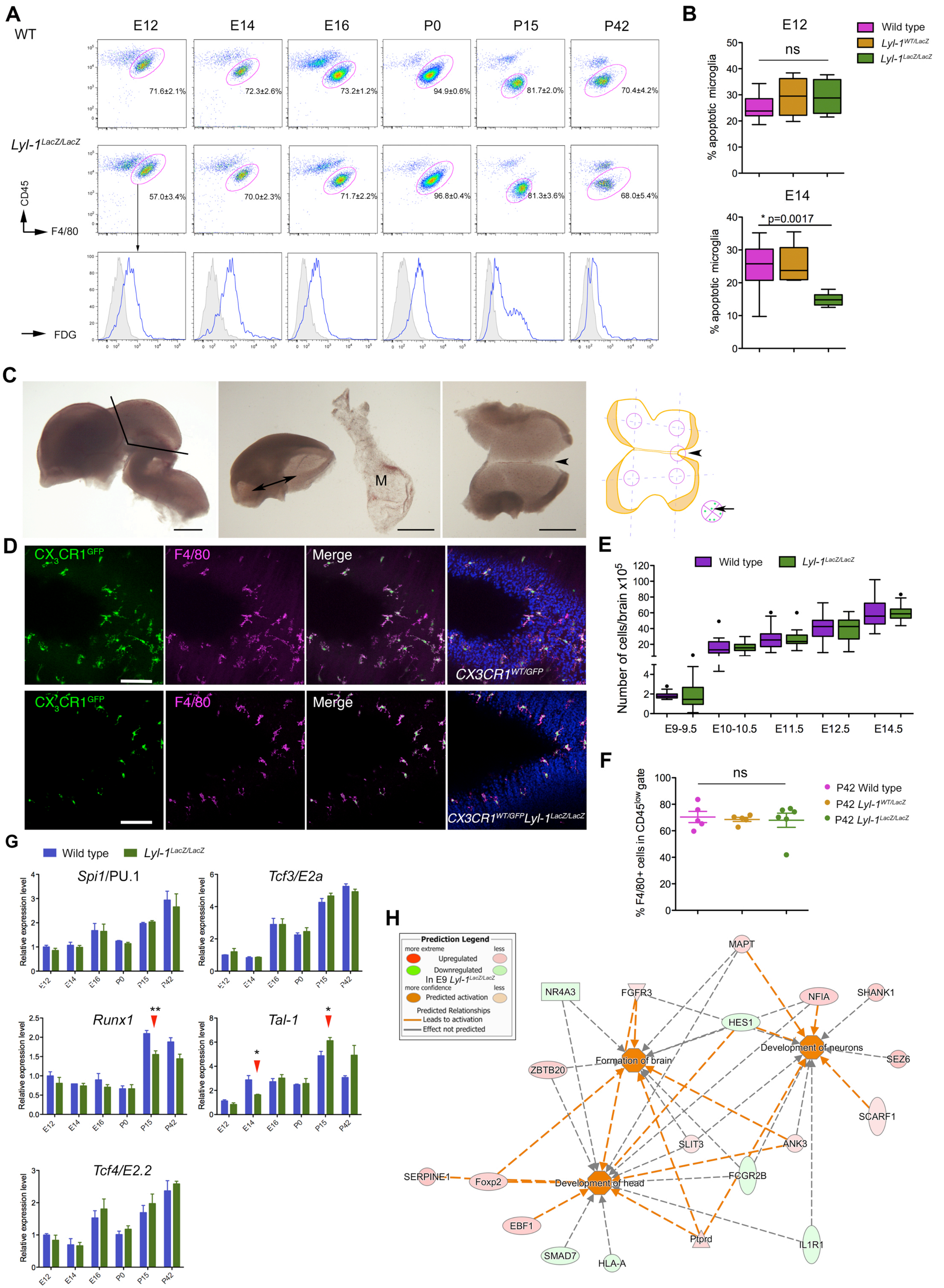

#### *Supplemental Tables*

##### *Supplemental Table 1: GSEA gene sets enriched in WT MΦP at E10*

Compared to E9 MΦP, E10 progenitors were enriched in pathways involved in the regulation of erythro-myeloid lineages (Green), in inflammatory signaling and in metabolism (pink).

| Rank | Gene sets enriched in E10 WT compared to E9 WT MΦP | Size | NES | Nom.p-value | FDR q-val |
|---|---|---|---|---|---|
| 1 | Pyruvate metabolism | 34 | -1.98 | 0.0 | 0.027 |
| 2 | Diurnally regulated genes with circadian orthologs | 40 | -1.93 | 0.0 | 0.039 |
| 3 | Butanoate metabolism | 16 | -1.90 | 0.0 | 0.040 |
| 4 | Epo signaling pathway | 15 | -1.88 | 0.0 | 0.045 |
| 5 | Oxidative stress response | 45 | -1.87 | 0.0 | 0.038 |
| 6 | ATM signaling pathway | 17 | -1.86 | 0.0032 | 0.034 |
| 7 | Blood coagulation | 37 | -1.86 | 0.0 | 0.030 |
| 8 | Toll receptor signaling pathway | 38 | -1.85 | 0.0 | 0.031 |
| 9 | Adipogenesis | 125 | -1.85 | 0.0 | 0.028 |
| 10 | Estrogen signaling pathway | 18 | -1.85 | 0.0 | 0.026 |
| 11 | TNF-R2 signaling pathway | 17 | -1.81 | 0.0 | 0.038 |
| 12 | DNA damage response only ATM dependent | 77 | -1.81 | 0.0028 | 0.036 |
| 13 | TGFβ signaling pathway | 109 | -1.80 | 0.0 | 0.034 |
| 14 | Alpha6 beta4 integrin | 69 | -1.80 | 0.0 | 0.036 |
| 15 | Propanoate metabolism | 23 | -1.78 | 0.0032 | 0.040 |
| 16 | TPO signaling pathway | 18 | -1.78 | 0.0 | 0.039 |
| 17 | Folate metabolism | 46 | -1.77 | 0.0029 | 0.040 |
| 18 | Gene regulation by peroxisome proliferators via ppara-alpha | 42 | -1.77 | 0.0 | 0.038 |
| 19 | Oxidative stress induced gene expression via NRF2 | 16 | -1.76 | 0.0033 | 0.039 |
| 20 | Steroids metabolism | 15 | -1.74 | 0.0016 | 0.047 |
| 21 | TGFβ signaling pathway (wp560) | 51 | -1.74 | 0.0013 | 0.045 |
| 22 | Il-3 signaling pathway | 39 | -1.74 | 0.0014 | 0.046 |
| 23 | Mets effect on macrophage differentiation | 15 | -1.73 | 0.006600 | 0.049 |

***Supplemental table 2*: GSEA pathways related to developmental patterning enriched in E9 Lyl-1$^{LacZ/LacZ}$ MΦP.**

| | GSEA Pathway up in E9 Mut vs WT *MΦP* | SIZE | NES | NOM p-val | FDR q-val | |
|---|---|---|---|---|---|---|
| 4 | Cadherin signaling pathway | 97 | 1.90 | 0.000 | 0.022 | *Cdh3; Cdh5; Wnt3; Pcdhb6; Erbb4; Wnt7b; Wnt1; Tcf7l1; Pcdh8; Cdh20; Wnt4; Cdh9; Pcdh19; Ctnna3; Wnt7a; Cdh8; Pcdha4; Wnt9a; Pcdh18; Wnt5a; Cdh6; Src; Cdh18; Erbb3; Cdh4; Erbb2; Cdh24; Fzd9; Celsr2; Fzd8; Pcdhb4; Cdh11; Celsr1; Acta2; Wnt2b; Wnt11; Pcdhb5; Cdh7; Fat3* |
| 12 | Cytoskeletal regulation by RHO GTPase | 60 | 1.57 | 0.013 | 0.242 | *Myh3; Myh1; Rhoj; Pak6; Myo3a; Rhoc; Myh6; Myh14; Myh2; Tubb6; Stmn4; Acta2; Myh7; Actg2; Pak3; Vasp; Pak4; Actbl2; Myh4; Arpc4; Rac2; Pak1* |
| 13 | Wnt signaling pathway | 225 | 1.55 | 0.002 | 0.247 | *Myh3; Myh1; Nkd1; Celsr3; Sfrp5; Cdh3; Cdhr1; Wnt3; Pcdhb6; Wnt7b; Wnt1; Tcf7l1; Pcdh8; Dkk4; Cdh20; Sfrp2; Wnt4; Cdh9; Myh6; Mycn; Pcdh19; Ctnna3; Axin2; Wnt7a; Cdh8; Pcdha4; Wnt9a; Pcdh18; Acvr1b; Myh2; Wnt5a; Prkcz; Cdh6; Smarcd3; Pygo1; Edn1; Tle2; Cdh18; Cdh4; Gng13; Cdh24; Arr3; Fzd9; Celsr2; Fzd8; Pcdhb4; Cdh11; Celsr1; Gng12; Gna11; En1; Ctnnal1; Acta2* |
| 25 | Integrin signaling pathway | 162 | 1.41 | 0.013 | 0.340 | *Itga2b; Lama4; Col17a1; Col11a2; Lamb2; Col8a2; Parvb; Itgae; Col18a1; Col19a1; Col5a3; Rhoc; Col9a3; Rnd2; Parva; Itga1; Col4a2; Rnd1; Col5a1; Col6a1; Pik3c2g; Grap; Col11a1; Col1a1; Col6a2; Col4a1; Col3a1; Src; Col1a2; Col12a1; Col9a1; Itgbl1; Itgb3; Mapk3; Lamc2; Col10a1; Col4a6; Lims2; Map2k3; Itga8; Col4a3* |

***Supplemental table 3: Lyl-1-positive MΦ$^{Prim}$ from E10-YS could play a role in heart development and function.*** KEGG gene sets down-regulated in E10 MΦP in *Lyl-1*$^{LacZ/LacZ}$, compared to WT.

| Rank | Gene sets | SIZE | NES | NOM p-val | FDR q-val |
|---|---|---|---|---|---|
| 1 | Myl3 pathway (Reactome mm) | 18 | -2.08 | 0.000 | 0.004 |
| 2 | Myl1 pathway (Reactome mm) | 20 | -2.08 | 0.000 | 0.003 |
| 3 | Tcap pathway (Reactome mm) (Titin, muscle) | 15 | -2.04 | 0.000 | 0.004 |
| 4 | Neb pathway (Reactome mm) (Nebulin, muscle, myopathy) | 16 | -2.00 | 0.000 | 0.006 |
| 5 | Myl4 pathway (Reactome mm) | 17 | -1.99 | 0.000 | 0.005 |
| 6 | Myh8 pathway (Reactome mm) | 21 | -1.94 | 0.000 | 0.011 |
| 7 | Myh6 pathway (Reactome mm) | 21 | -1.93 | 0.000 | 0.012 |
| 8 | Myh3 pathway (Reactome mm) | 22 | -1.92 | 0.000 | 0.012 |
| 9 | Mybpc3 pathway (Reactome mm) | 23 | -1.87 | 0.000 | 0.025 |
| 10 | Mybpc2 pathway (Reactome mm) | 23 | -1.87 | 0.000 | 0.023 |
| 11 | Mybpc1 pathway (Reactome mm) | 23 | -1.87 | 0.000 | 0.021 |
| 12 | Myl2 pathway (Reactome mm) | 28 | -1.82 | 0.000 | 0.047 |
| 13 | Desmin pathway (Reactome mm) (Muscle-cardiopathy) | 27 | -1.77 | 0.000 | 0.092 |
| 14 | Antigen processing and presentation (Kegg mm) | 34 | -1.65 | 0.005 | 0.386 |
| 15 | Ncor1 pathway (Reactome mm) | 44 | -1.65 | 0.006 | 0.364 |
| 16 | Dmd (dystrophin) pathway (Reactome mm) | 51 | -1.62 | 0.009 | 0.442 |
| 17 | Hmgb1 pathway (Reactome mm) | 24 | -1.61 | 0.016 | 0.482 |
| 18 | Ap2m1 pathway (Reactome mm) | 24 | -1.61 | 0.003 | 0.468 |
| 19 | Fgf1_pathway (Reactome mm) | 30 | -1.60 | 0.011 | 0.478 |
| 20 | Cytokine-cytokine receptor interaction (Kegg mm) | 199 | -1.58 | 0.002 | 0.535 |

*Supplemental Table 4: GSEA transcription factors*

**a.** Differential expression of transcription factor gene sets between E9 and E10 MΦP in a WT context.

| Rank | Gene set enriched in WT MΦP at E9 | SIZE | NES | NOM p-val | FDR q-val |
|---|---|---|---|---|---|
| 1 | REST (TFACTS-MM) | 19 | 1.68 | 0.010 | 0.182 |
| **Rank** | **Gene sets enriched in WT MΦP at E10** | **SIZE** | **NES** | **NOM p-val** | **FDR q-val** |
| 1 | GATA1 (TFACTS-MM) | 23 | -2.53 | 0.000 | 0.000 |
| 2 | ESR1 (TFACTS-MM) | 47 | -1.95 | 0.000 | 0.015 |
| 3 | JUN (TF-MM-Zhao) | 93 | -1.94 | 0.000 | 0.011 |
| 4 | JUND (TFACTS-MM) | 24 | -1.92 | 0.000 | 0.011 |
| 5 | CEBPB (TF-MM-Zhao) | 20 | -1.92 | 0.000 | 0.009 |
| 6 | RARA (TFACTS-MM) | 51 | -1.91 | 0.000 | 0.008 |
| 7 | JUN (TFACTS-MM) | 103 | -1.90 | 0.000 | 0.009 |
| 8 | SREBF1 (TFACTS-MM) | 44 | -1.90 | 0.000 | 0.008 |
| 9 | JUNB (TFACTS-MM) | 15 | -1.89 | 0.000 | 0.007 |
| 10 | SFPI1 (TF-MM-Zhao) | 41 | -1.89 | 0.000 | 0.007 |
| 11 | SMAD7 (TFACTS-MM) | 15 | -1.88 | 0.000 | 0.008 |
| 12 | SREBF2 (TFACTS-MM) | 31 | -1.87 | 0.001 | 0.008 |
| 13 | LEF1 (TFACTS-MM) | 43 | -1.84 | 0.001 | 0.011 |
| 14 | YY1 (TFACTS-MM) | 37 | -1.84 | 0.002 | 0.011 |
| 15 | SP1 (TFACTS-MM) | 352 | -1.82 | 0.000 | 0.012 |
| 16 | SMAD4 (TFACTS-MM) | 38 | -1.82 | 0.005 | 0.012 |
| 17 | ETS1 (TFACTS-MM) | 114 | -1.79 | 0.001 | 0.014 |
| 18 | ELK1 (TFACTS-MM) | 15 | -1.79 | 0.000 | 0.014 |
| 19 | AR (TFACTS-MM) | 45 | -1.78 | 0.001 | 0.015 |
| 20 | SMAD3 (TFACTS-MM) | 53 | -1.78 | 0.000 | 0.014 |
| 21 | ATF2 (TFACTS-MM) | 28 | -1.75 | 0.005 | 0.020 |
| 22 | FOXO1 (TFACTS-MM) | 135 | -1.74 | 0.000 | 0.021 |
| 23 | HOXC8 (TF-MM-Zhao) | 25 | -1.72 | 0.008 | 0.025 |
| 24 | RXRA (TFACTS-MM) | 20 | -1.72 | 0.003 | 0.024 |

**b.** Transcription factor gene sets modified in E9 *Lyl-1*$^{LacZ/LacZ}$ MΦP, compared to WT.

| Rank | Gene set enriched in E9 *Lyl-1*$^{LacZ/LacZ}$ MΦP | SIZE | NES | NOM p-val | FDR q-val |
|---|---|---|---|---|---|
| 1 | SMAD1 (TF-MM-Zhao) | 15 | 1.66 | 0.024 | 0.363 |
| **Rank** | **Gene sets depleted in E9 *Lyl-1*$^{LacZ/LacZ}$ MΦP** | **SIZE** | **NES** | **NOM p-val** | **FDR q-val** |
| 1 | SFPI1 (TF-MM) | 41 | -0.74 | 0.000 | 0.000 |
| 2 | SPI1 (TFACTS-MM) | 73 | -0.58 | 0.000 | 0.000 |
| 3 | STAT1 (TFACTS-MM) | 42 | -0.62 | 0.000 | 0.000 |
| 4 | JUN (TF-MM-Zhao) | 93 | -0.49 | 0.000 | 0.004 |
| 5 | PPARG (TF-MM-Zhao) | 37 | -0.56 | 0.000 | 0.019 |
| 6 | TBP (TF-MM-Friard) | 39 | -0.53 | 0.000 | 0.049 |
| 7 | SFPI1 (TF-MM) | 29 | -0.56 | 0.000 | 0.043 |
| 8 | JUN (TFACTS-MM) | 103 | -0.43 | 0.000 | 0.046 |
| 9 | NFKB1 (TF-MM-Zhao) | 65 | -0.46 | 0.002 | 0.047 |
| 10 | NFKB1 (TFACTS-MM) | 111 | -0.42 | 0.000 | 0.045 |
| 11 | CEBPA (TFACTS-MM) | 93 | -0.43 | 0.000 | 0.046 |

***Supplemental table 5: Cytokines used in this study***

| ***Cytokine*** | ***Concentration*** | ***Supplier*** |
|---|---|---|
| Murine recombinant Stem Cell factor | 50ng/mL | Peprotech; Ref: 315-03 |
| Human recombinant EPO | 3U/mL | In house production |
| Murine recombinant IL-3 | 10ng/mL | Peprotech; Ref: 213-13 |
| Human recombinant IL-6 | 10ng/mL | A gift from Sam Burstein, Maryville, USA |
| Human recombinant CSF-1 | 10ng/mL | Peprotech; Ref: 300-25 |
| Human recombinant TPO | 10ng/mL | A gift from Kirin Brewery, Tokyo, Japan |

*Supplemental table 6: Antibodies and fluorescent stains used throughout this study*

| ***Antibody name*** | ***Clone*** | ***Supplier/Reference*** | ***Fluorochrome/ Chromogen*** |
|---|---|---|---|
| Ter119 | TER-119 | Biolegend; 116205<br>Biolegend; 116208<br>eBioscience 17-5921-82 | FITC<br>PE<br>APC |
| F4/80 | BM8<br>CI:A3-1<br>BM8<br>BM8 | eBioscience; 53-4801-82<br>Biolegend; 122606<br>eBioscience; 12-4801-82<br>Biolegend ; 123116 | Alexa fluor 488<br>FITC<br>PE<br>APC |
| GR-1 | RB6-8C5<br>RB6-8C5<br>RB6-8C5 | BD-Pharmingen; 553127<br>Biolegend; 108416<br>Biolegend , 108437 | FITC<br>PE-Cy7<br>BV510 |
| CD45 | 30-F11<br>30-F11<br>30-F11<br>30-F11 | BD-Pharmingen; 553081<br>BD-Pharmingen; 553082<br>eBioscience; 25-0451-82<br>Biolegend; 103124 | PE<br>PE-Cy5<br>PE-Cy7<br>Alexa fluor 647 |
| CD31 | MEC13.3<br>390<br>MEC13.3<br>MEC13.3<br>MEC13.3 | Biolegend; 102514<br>Biolegend; 102408<br>Biolegend; 102419<br>Biolegend; 102516<br>BD-Pharmingen; 583089 | Alexa fluor 488<br>PE<br>PE-Cy5.5<br>Alexa fluor 647<br>BV510 |
| Kit | 2B8<br>2B8<br>2B8 | Biolegend; 05824<br>BD-Pharmingen; 553358<br>Biolegend ; 105825 | Alexa fluor 488<br>APC<br>APC-Cy7 |
| CD11b | M1/70 | eBioscience; 45-0112-82<br>Biolegend; 101215<br>Biolegend; 101212<br>eBioscience; 47-0112-82 | PE-Cy5.5<br>PE-Cy7<br>APC<br>APC-eFluor 780 |
| Sca-1 | D7 | Biolegend; 108113 | PE-Cy7 |
| MHC-II | M5/114.15.2 | Biolegend; 107607 | PE |
| Annexin V | NA | Biolegend; B206040 | FITC |
| Anti-Brdu | B44 | BD-Pharmingen; 51-23619L | APC |
| 7AAD | NA | BD-Pharmingen; 51-68981E | Dye |
| FDG | NA | Molecular probe; Thermo Fisher; F1179 | Dye |

*Supplemental table 7: Primers used in this study*

| ***Gene*** | | ***Sequence*** |
|---|---|---|
| ***Actin*** | Primer forward: | 5′- CTTCTTTGCAGCTCCTTCGT-3′ |
| | Primer reverse: | 5′- ATCACACCCTGGTGCCTAG-3′ |
| ***Hprt*** | Primer forward: | 5′- TGATTATGGACAGGACTGAAAGA-3′ |
| | Primer reverse: | 5′- AGCAGGTCAGCAAAGAACTTATAG-3′ |
| | Primer reverse: | 5'-TGGTTCTGCATCGACTTCTG-3′ |
| ***Tubulin*** | Primer forward: | 5′- GGGTGGAGGCACTGGCT-3′ |
| | Primer reverse: | 5′- GTCAGGATATTCTTCCCGGATCT-3′ |
| ***c-Maf*** | Primer forward: | 5'- CTGCCGCTTCAAGAGGGTGCAGC -3' |
| | Primer reverse: | 5'- GATCTCCTGCTTGAGGTGGTC -3' |
| ***Lyl-1*** | Primer forward: | 5'- TGGACTGACAAACCTGACCA-3' |
| | Primer reverse: | 5'- TGGACCCCACGGATAGAATG-3' |
| ***Tal-1/Scl*** | Primer forward: | 5'- CATGTTCACCAACAACAACCG-3' |
| | Primer reverse: | 5'- GGTGTGAGGACCATCAGAAATCTC-3' |
| ***Runx1*** | Primer forward: | 5'- CTCCGTGCTACCCACTCACT-3' |
| | Primer reverse: | 5'- ATGACGGTGACCAGAGTGC-3' |
| ***Meis1*** | Primer forward: | 5'-TGAAGTAGGAAGGGAGCCAG-3′ |
| | Primer reverse: | 5'-GCCTACTCCATCCATACCCC-3′ |
| ***Lmo2*** | Primer forward: | 5'-ATGTCCTCGGCCATCGAAAG-3′ |
| | Primer reverse: | 5'-CGGTCCCCTATGTTCTGCTG-3′ |
| ***PU.1*** | Primer forward: | 5'- GCTATACCAACGTCCAATGCA-3' |
| | Primer reverse: | 5'-TGTGCGGAGAAATCCCAGTA-3' |
| ***Tcf3/E2A*** | Primer forward: | 5'-GGGCTCTGACAAGGAACTGA-3′ |
| | Primer reverse: | 5'-AGTCCTGAGCCTGCAAACTG-3′ |
| ***Tcf4/E2.2*** | Primer forward: | 5'-GCCTCTTCACAGTAGTGCCAT-3′ |
| | Primer reverse: | 5'-TCCCTGTTGTAGTCGGCAGT-3′ |
| ***Csfr1*** | Primer forward: | 5'-GCAGTACCACCATCCACTTGTA-3' |
| | Primer reverse: | 5'-GTGAGACACTGTCCTTCAGTGC-3' |
| ***Cx3cr1*** | Primer forward: | 5′ - AGTTCCCTTCCCATCTGCTC-3′ |
| | Primer reverse: | 5′ - GCCACAATGTCGCCCAAATA-3′ |
| ***Irf8*** | Primer forward: | 5'- CGGGGCTGATCTGGGAAAAT-3' |
| | Primer reverse: | 5'- CACAGCGTAACCTCGTCTTC-3' |
| ***Myb*** | Primer forward: | 5′-AGCGTCACTTGGGGAAAACT-3′ |
| | Primer reverse: | 5′- AGTCGTCTGTTCCGTTCTGT-3′ |
